# Activation of NLRP3 Inflammasome by Virus-like Particles of Human Polyomaviruses in Macrophages

**DOI:** 10.1101/2021.11.15.468577

**Authors:** Asta Lučiūnaitė, Indrė Dalgėdienė, Rapolas Žilionis, Kristina Mašalaitė, Milda Norkienė, Andrius Šinkūnas, Alma Gedvilaitė, Indrė Kučinskaitė-Kodzė, Aurelija Žvirblienė

## Abstract

Viral antigens can activate phagocytes inducing inflammation but the mechanisms are barely explored. This study aimed to investigate the capability of viral oligomeric proteins of different structure to induce inflammatory response in macrophages. Human THP-1 cell line was used to prepare macrophages which were treated with filamentous nucleocapsid-like particles (NLPs) of paramyxoviruses and spherical virus-like particles (VLPs) of human polyomaviruses. The effects of viral proteins on cell viability, pro-inflammatory cytokines’ production and formation of NLRP3 inflammasome components, ASC specks, were investigated. Filamentous NLPs did not induce inflammation markers while spherical VLPs mediated inflammatory response followed by NLRP3 inflammasome activation. Inhibitors of cathepsins and K^+^ efflux decreased IL-1β levels and cell death indicating a complex inflammasome activation process. Similar activation pattern was observed in primary human macrophages treated with VLPs. Single cell RNAseq analysis of THP-1 cells revealed several cell activation states characterized by high expression of inflammation-related genes. This study provides new insights into interaction of viral proteins with innate immune cells and suggests that structural properties of oligomeric proteins may define cell activation pathways.

## Introduction

The components of innate immunity such as macrophages play a key role in the onset and progression of inflammatory and age-related diseases (Oishi & Manabe, 2016; Parisi *et al*, 2018). Macrophages are considered as a potential target for treatment of many diseases, therefore, molecular mechanisms related to their activation is currently at the top of research (Zhang *et al*, 2021). Besides, macrophages are known to recognize the structural properties of the activation agents and influence the inflammatory response via inflammasome activation (Rabolli *et al*, 2016a; Shu & Shi, 2018). Inflammasomes are intracellular protein complexes representing important components of the innate immune system (de Alba, 2019). The best described representative is NLRP3 inflammasome. It contains three major components – nucleotide-binding and oligomerization domain-like receptor, apoptosis-associated speck-like protein containing CARD (ASC) and pro-caspase-1. NLRP3 inflammasome assembly results in IL-1β release and inflammatory cell death – pyroptosis (Swanson *et al*, 2019). Endogenous and external factors can trigger its assembly. Its assembly can be triggered by endogenous and external factors. Activation of NLRP3 inflammasome is associated with various diseases, including gout and Alzheimer’s disease (Fusco *et al*, 2020). Our previous study showed that NLRP3 inflammasome is activated by amyloid beta (Aβ), both oligomers and protofibrils (Luciunaite *et al*). Inflammasome activation was also induced by α-synuclein (Codolo *et al*, 2013), and tau oligomers (Ising *et al*, 2019). However, the most explored inflammasome triggers are polymeric nanoparticles (Doshi & Mitragotri, 2010), cholesterol crystals (Samstad *et al*, 2014), airborne pollutants such as silica (Dostert *et al*, 2008). The structural properties of synthetic nanoparticles determine the outcome of cell activation (Baranov *et al*, 2020). Previously, we have demonstrated that spherical oligomeric proteins of viral origin induce inflammatory responses in macrophages but the mechanism and molecular components of this process are not yet confirmed (Dalgediene *et al*, 2018). In addition, another study demonstrated that the nucleocapsid (N) protein of Zika virus activated the inflammasome (Wang *et al*, 2018). Repetitive and latent viral infections are potential agents of inflammation (Maloney *et al*, 2013). This frame of reference draws attention to the latent viral infections and their role in inflammation.

It is well known that some viruses lie dormant within the host after an acute infection. According to World Health Organisation, 50-80% of world population is infected by polyomaviruses (PyVs) in the childhood (De Gascun & Carr, 2013). The effects of these viruses and their antigens on innate immunity throughout the life are not fully understood. For example, herpes virus in its latent form can cause multisymptom illness with a broad range of simultaneous symptoms, such as cognitive disorders, depression, fatigue (Maloney *et al*., 2013). However, the mechanism of multisymptomatic illness is unclear. Investigation of inflammatory responses induced by proteins of viruses causing acute and latent infection may indicate whether these viral proteins alone are capable of activating the immune system. Moreover, VLPs are already established as carriers and adjuvants in vaccines. Components of vaccines can activate the inflammasome as it was reported for the ISCOMATRIX adjuvant (Wilson *et al*, 2014). Studying how different viral proteins especially VLPs stimulate the immune cells would allow the selection of better tools for vaccination.

Inflammatory reactions induced by synthetic polymeric particles in macrophages vary depending on their size and shape (Shu & Shi, 2018; Rabolli, 2016). Similarly, viral proteins may also fate cellular response depending on their structural properties. Our study was aimed to investigate how recombinant viral proteins induce inflammatory response in macrophages focusing on inflammasome activation. The VLPs derived from major capsid protein VP1 of human PyVs – Karolinska Institute (KI) PyV and Merkel cell (MC) PyV, self-assembling to spherical particles about 20-60 nm in diameter (Norkiene *et al*, 2015b), were selected as typical representatives of spherical oligomeric proteins. The NLPs of measles and mumps viruses were chosen as a model of oligomeric proteins forming filamentous rod-shaped structures (Samuel *et al*, 2003; Zvirbliene *et al*, 2007). We showed that inflammatory responses and activation of NLRP3 inflammasome in macrophages depend on the structural properties of viral proteins. This study provides new insights on the ability of multimeric viral antigens to induce inflammatory response in innate immune cells.

## Results

We investigated the ability of viral multimeric proteins to induce inflammatory response in macrophages focusing on inflammasome activation. For this study, we have selected structurally diverse viral proteins: filamentous NLPs of measles and mumps viruses (*Paramyxoviridae* family) that usually cause acute infections, and spherical VLPs derived from VP1 of PyVs (*Polyomaviridae* family) that generally induce latent viral infections. We also investigated whether viral oligomeric proteins of diverse structure can determine different patterns of cell activation.

### NLPs did not activate THP-1 macrophages despite their interaction with the cells

First at all, we investigated the uptake of viral proteins by THP-1 macrophages. The cells were treated with recombinant NLPs of measles and mumps viruses, as well as recombinant VLPs of PyVs, for 24 h to observe the uptake of multimeric viral proteins. The NLPs and VLPs were immunostained with the respective monoclonal antibodies. We also immunostained the cells for the macrophage and lysosomal marker CD68. The uptake of both NLPs and VLPs was detected microscopically demonstrating their interaction with THP-1 macrophages (Fig 1).

**Figure 1.**
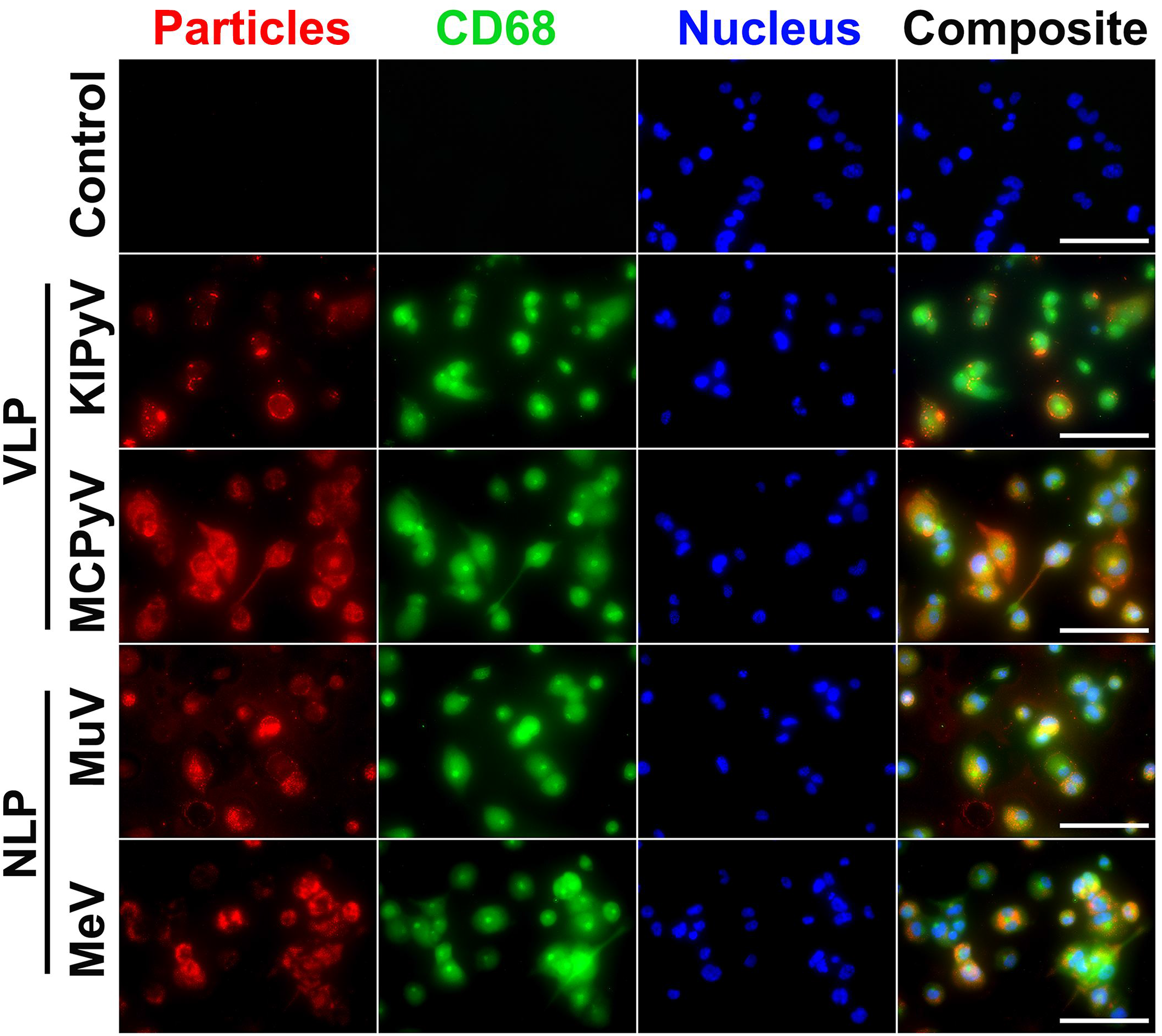
The uptake of NLPs of measles and mumps viruses, and PyV-derived VLPs by THP-1 macrophages. THP-1 macrophages were treated with recombinant viral proteins for 24 h at 20 µg/ml concentration. Cells were immunostained with anti-NLP and anti-VLP monoclonal antibodies (red), anti-CD68 – macrophage and lysosomal marker (green), nuclear stain Hoechst33342 (blue) and analysed by fluorescence microscopy. The negative control – secondary antibody alone is referred as a control. Images were taken using 40× objective. The scale bars indicate 100 μm. All experiments were performed in triplicate. Representative images of one experiment are shown.

To investigate macrophage activation by multimeric viral proteins, we started with recombinant NLPs of measles and mumps viruses forming filamentous structures. These NLPs are long rod-shaped structures, about 20 nm in diameter, mimicking the nucleocapsids of native viruses (Samuel *et al*, 2002a; Slibinskas *et al*, 2004). Treatment of THP-1 macrophages with recombinant NLPs did not cause any inflammatory response according to TNF-α release data (Fig 2A). The NLPs also did not activate the inflammasome as no change in IL-1β (Fig 2B) and lactate dehydrogenase (LDH) release (Fig 2C) was detected. In addition, the investigated viral proteins were not cytotoxic according to propidium iodide (PI) and Hoechst nuclear staining assay (Fig 2D and E). We concluded that recombinant NLPs did not induce the inflammatory response in THP-1 macrophages.

**Figure 2.**
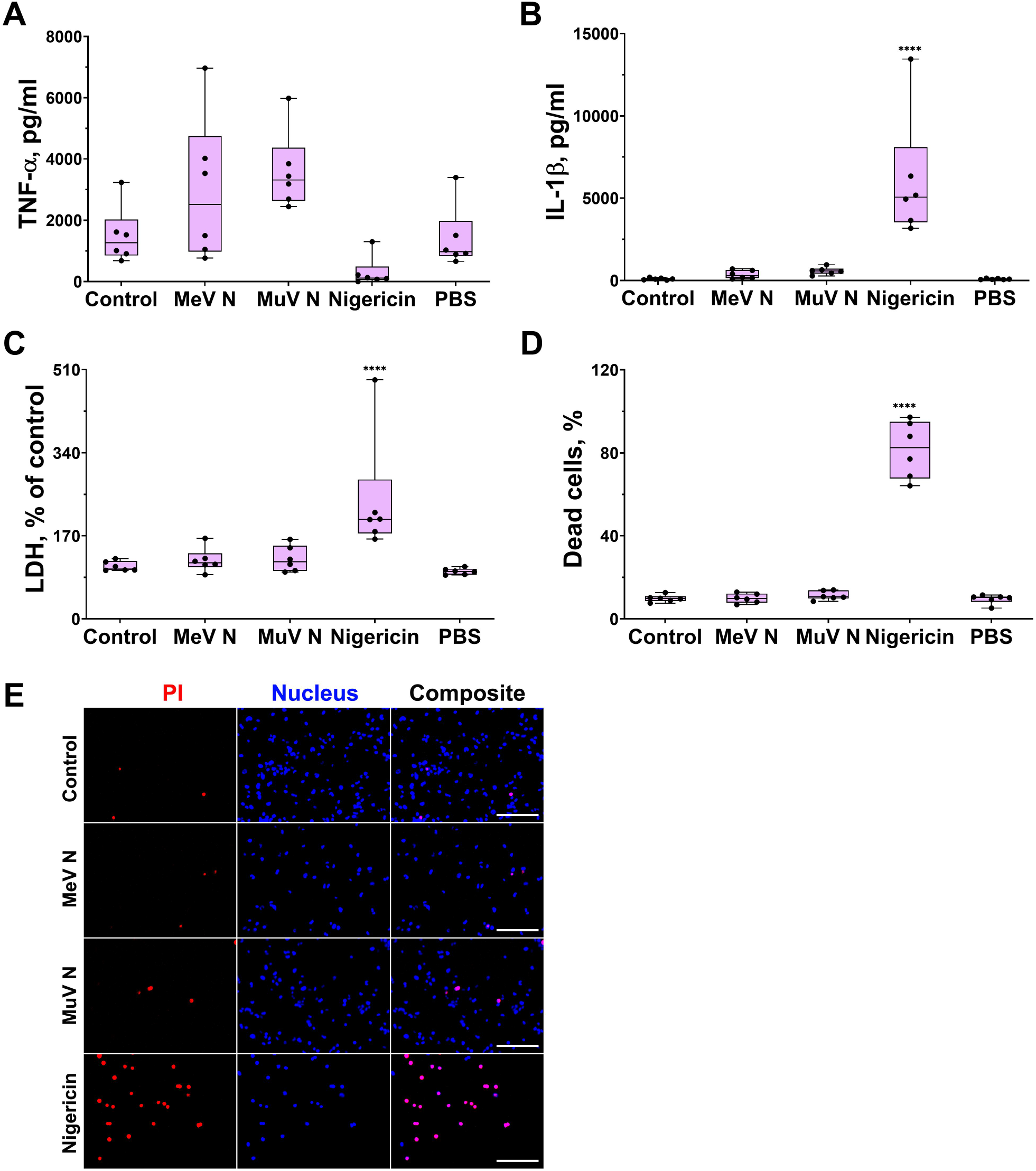
NLPs of measles and mumps viruses did not activate THP-1 macrophages. A, B Macrophages were treated with viral proteins (20 µg/ml) for 24 h. Nigericin (10 μM) was used as a positive control. (A) TNF-α and (B) IL-1β secretion determined by ELISA. C, D Cytotoxicity assessed by (C) LDH assay and (D) PI and Hoechst nuclear staining. PI indicates dead cells and Hoechst stains all cell nuclei. (E) Representation of the staining. Images were taken using 20× objective. The scale bars indicate 200 μm. Data information: Data are represented using box plots, *p < 0.05, **p < 0.01, ***p < 0.001, ****p < 0.0001, one-way ANOVA followed by Tukey’s multiple comparison test, stars show statistically significant results compared to control.

### PyV-derived VLPs induced inflammatory response followed by NLRP3 inflammasome activation in THP-1 macrophages

Next, we investigated macrophage activation by PyV-derived VLPs of spherical structures that are similar to native viruses in their shape and size. In order to evaluate the effects of diverse VLPs we have selected PyV-derived VLPs of different sizes, ranging 20-60 nm in diameter (Norkiene *et al*, 2015a). KIPyV-derived VP1 proteins form heterogeneous particles, ranging from 20 to 60 nm, while MCPyV-derived VP1 proteins form more homogenous VLPs, 45-50 nm in diameter. Thus, we investigated VLPs of different sizes. Treatment of human THP-1 macrophages with PyV-derived VLPs induced cell activation and inflammatory response according to TNF-α and IL-6 release (Fig 3A and B). Therefore, we assumed that VLPs activated the NFκ-β signalling pathway, which could be a priming step for further inflammasome activation. In addition, we assayed the cells for anti-inflammatory cytokine IL-10 secretion and did not detect its release after VLP treatment (detected IL-10 values were below the assay detection limit, so, equalised 0). PyV-derived VLPs induced not only secretion of inflammatory cytokines but also provoked cell death as LDH release was increased (Fig 3D) and dead cells were detected using PI staining (Fig 3E). We also demonstrated that VLPs of different sizes induced cell activation signal of different strength. Large-sized (45-50 nm in diameter) homogenous VLPs of MCPyV induced higher inflammatory response as compared to heterogeneous KIPyV-derived VLPs (Fig 3).

**Figure 3.**
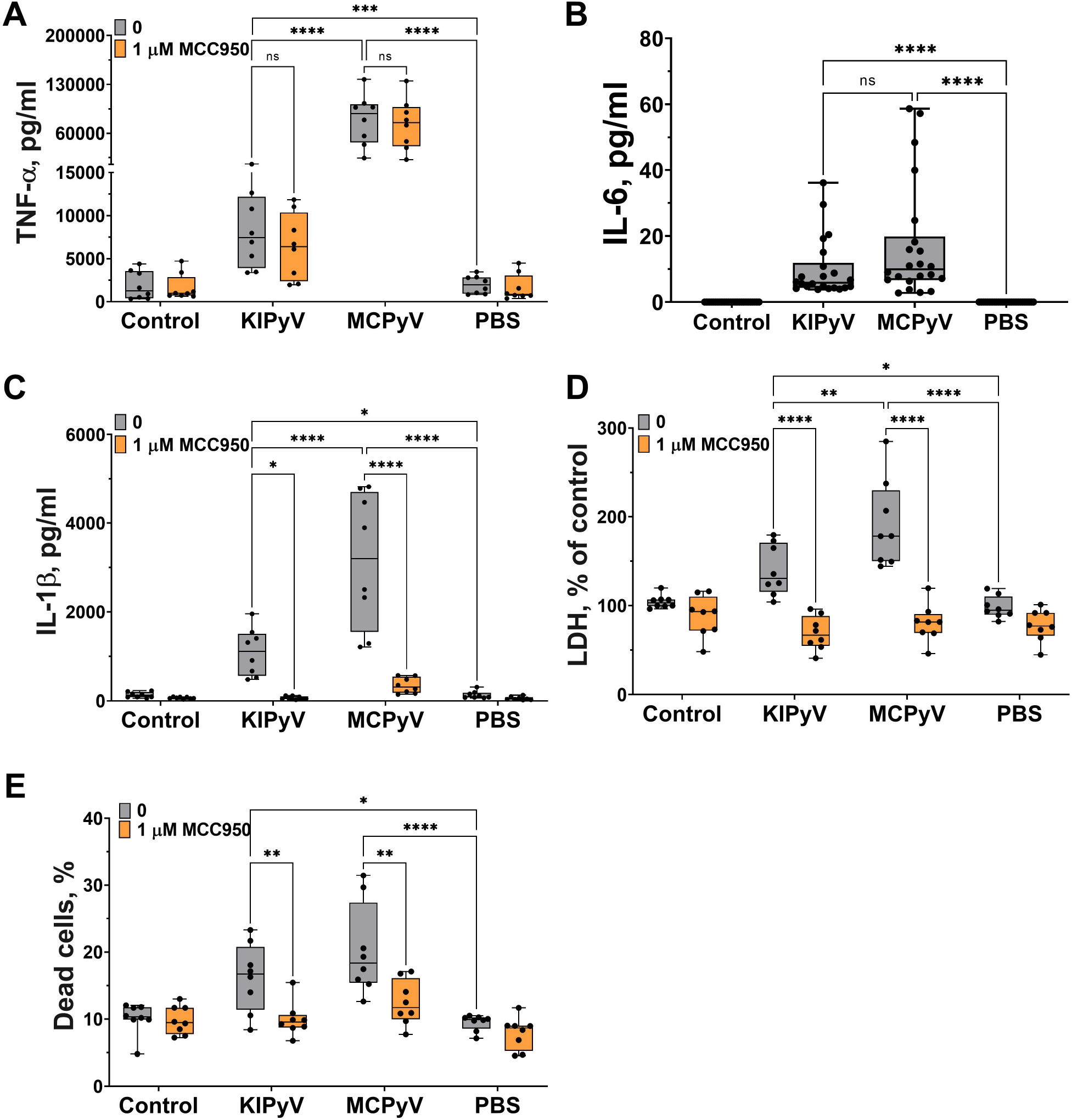
PyV-derived VLPs induced release of inflammatory cytokines TNF-α and IL-6 and activated NLRP3 inflammasome in human THP-1 macrophages. A-C Macrophages were treated with PyV-derived VLPs (20 µg/ml) for 24 h. Inhibitor MCC950 (1 μM) was added 30 min before treatment. (A) TNF-α, (B) IL-6, and (C) IL-1β secretion determined by ELISA. D, E Cytotoxicity assessed by (D) LDH assay and (E) PI and Hoechst nuclear staining. PI indicates dead cells (red) and Hoechst stains all cell nuclei (blue). Data information: Data are represented using box plots with dots showing independent experiments, *p < 0.05, **p < 0.01, ***p < 0.001, ****p < 0.0001, one-way ANOVA followed by Tukey’s multiple comparison test.

Then, we assessed whether PyV-derived VLPs induce activation of NLRP3 inflammasome in THP-1 macrophages. To test this, we used a small molecule MCC950 which specifically inhibits NLRP3 inflammasome. PyV-derived VLPs induced cell death (Fig 3D-F) and IL-1β release showing inflammasome activation (Fig 3C). Pre-treatment with MCC950 inhibitor before adding the VLPs significantly decreased this activation signal (Fig 3C-F). The inhibitor MCC950 did not influence the secretion level of other inflammatory cytokine TNF-α showing the specificity of inhibitor to inflammasome activation (Fig 3A). These data demonstrate that PyV-derived VLPs are potent inflammatory agents triggering inflammatory cytokine secretion and pyroptotic cell death.

In the next step, we investigated the formation of ASC specks that would indicate the assembly of the inflammasome. To detect ASC specks, we used THP-1-ASC-GFP macrophages. We identified ASC specks after treating cells with PyV-derived VLPs (Fig 4). However, MCC950 did not fully inhibit ASC speck formation (Fig 4B), which might be explained by ASC speck release from the inflammasome-activated cells. Secreted ASC specks can promote further maturation of inflammatory cytokines (Franklin *et al*, 2018). In addition, ASC specks could induce a subsequent inflammatory response by activating the surrounding cells.

**Figure 4.**
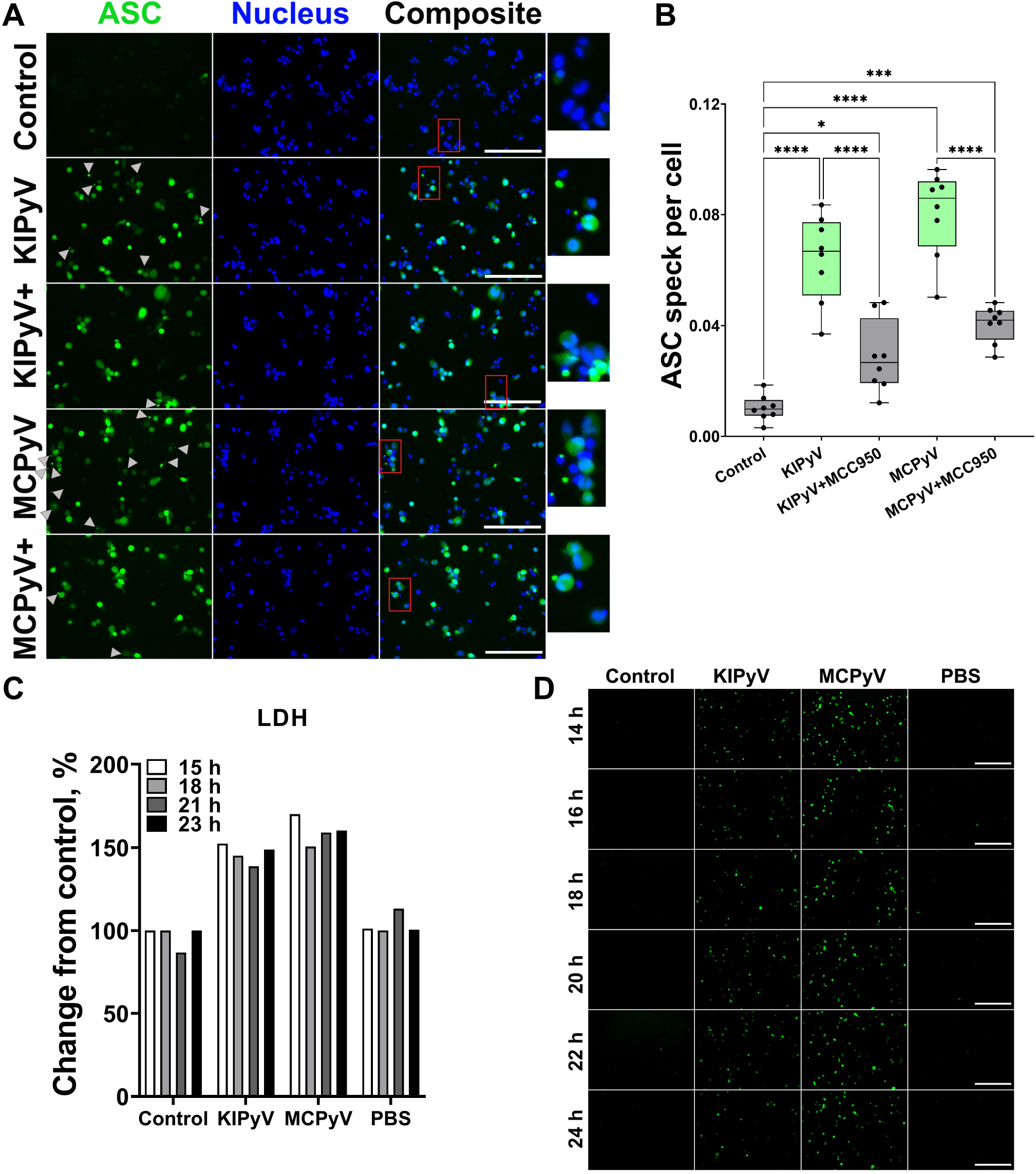
PyV-derived VLPs induced ASC speck formation in human THP-1 macrophages. A, B THP-1-ASC-GFP macrophages were treated for 24 h with PyV-derived VLPs (20 µg/ml), inhibitor MCC950 (1 μM) was added 30 min before treatment. Formation of ASC specks was visualised by fluorescent microscope. (A) Representative images of one experiment. Grey arrows show ASC specks. Red rectangles show magnified parts. “+” – refers to MCC950 pre-treatment. The images were taken using 20× objective. (B) Quantification of ASC speck count per cell. C, D Time lapse of PyV VLP-induced cell activation. (C) LDH assay was performed of cell culture supernatant in wilde-type THP-1 macrophages. (D) The formation of ASC specks detected in THP-1-ASC-GFP. Representative images of one experiment are shown. N = 1. Data information: ASC images were taken using 20× objective. The scale bars indicate 200 μm. For (B) data are represented using box plots with dots showing independent experiments, *p < 0.05, **p < 0.01, ***p < 0.001, ****p < 0.0001, one-way ANOVA followed by Tukey’s multiple comparison test. For (C) and (D) N = 1.

Next, we performed time-lapse experiments to observe the dynamics of cell activation by PyV-derived VLPs. We measured LDH release in wild-type THP-1 cells treated with PyV-derived VLPs for 15-23 h. There were no differences in LDH release within this time interval (Fig 4C). We also took the fluorescent microscopy images of THP-1-ASC-GFP after 14-24 h treatment with VLPs. Again, we did not find considerable differences within the 14- 24 h treatment period (Fig 4D). All cells were similar according to fluorescence intensity. We concluded that THP-1 macrophages are activated at about 15 h after VLP addition and their activation pattern remains constant from this time point.

To prove VLP-induced activation of the inflammasome, we assayed for active caspase-1, which is a major component of the inflammasome cascade, converting pro-IL-1β to its mature form. Having the time lapse experiment data, we collected cell culture supernatant for caspase-1 assay after 15 h – from the steady cell activation time point in order not to lose the active enzyme. The cleaved caspase-1 p20 fragment was immunodetected by WB after THP-1 treatment with VLPs (Fig 5A). NLRP3 inhibitor MCC950 reduced generation of the activated caspase-1. Then, we analysed caspase-1 activation at a single-cell level using FLICA reagent. It contains FAM-YVAD-FMK caspase-1 inhibitor probe, which covalently binds to only activated caspase-1. We revealed caspase-1 activation in some THP-1 macrophages after treatment with PyV-derived VLPs (Fig 5B). Large-sized MCPyV-derived VLPs induced a stronger response as compared to KIPyV-derived VLPs. Inflammasome inhibitor MCC950 reduced the level of activated caspase-1. We also stained VLP-treated cell cultures with PI to identify dead cells. Activated caspase-1 co-localised with dead cells confirming the pyroptotic cell death. Therefore, we concluded that PyV-derived VLPs induce inflammasome activation.

**Figure 5.**
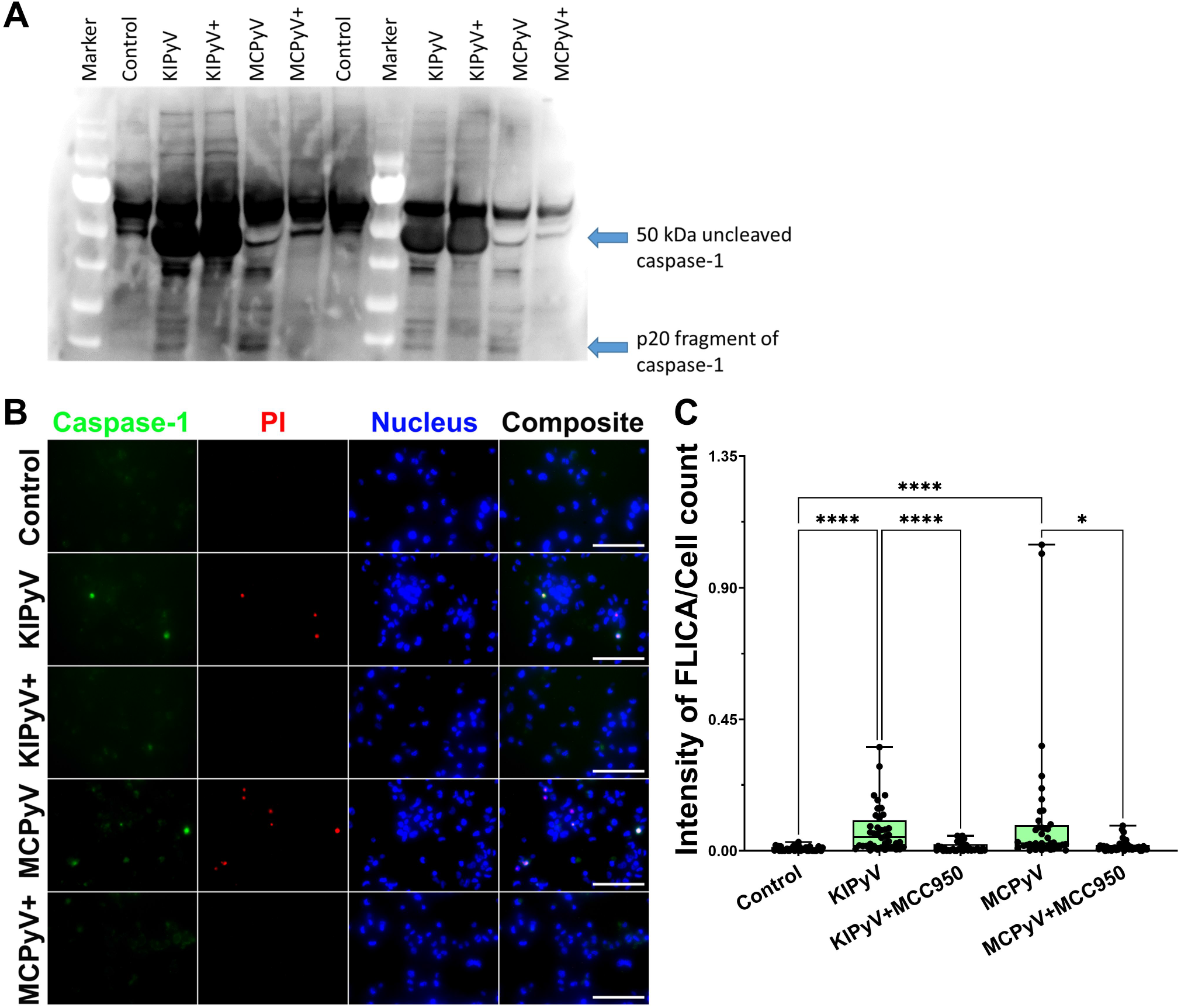
PyV-derived VLPs induced caspase-1 activation in human THP-1 macrophages. A–C Macrophages were treated for 15 h with PyV-derived VLPs (20 µg/ml), inhibitor MCC950 (1 μM) was added 30 min before treatment. (A) Cleaved caspase-1 was determined in cell supernatants by WB. Caspase-1: 50 kDa – pro-caspase-1; 20 kDa – cleaved caspase-1 (p20). “+” – refers to MCC950 pre-treatment. Duplicates of one experiment are shown. (A) Representative images of the activated caspase-1 staining by FLICA (green) reagent. Dead cell nuclear stain PI (red) and nuclear stain Hoechst (blue) was used. The images were taken using 20× objective. The scale bars indicate 100 μm. (C) Caspase-1 quantification according to FLICA analysis, n = 40 photos per condition. Data information: N = 1 independent experiment. For (C) data are represented using box- plots showing number of photos and analysed by Kruskal–Wallis test with Dunn’s post hoc, *p < 0.05, **p < 0.01, ***p < 0.001, ****p < 0.0001.

### The mechanism of VLP-mediated inflammasome activation is related to lysosomal damage

We further investigated the mechanism of inflammasome activation by PyV-derived VLPs. It is likely that accumulation of phagocytosed VLPs in lysosomes can damage them and induce the release of cathepsins, in particular cathepsin B (CtsB) that is one of the activators of NLRP3 inflammasome. We observed a significant decrease of IL-1β secretion after treatment of THP-1 cells with CtsB inhibitor Ca-074 Me (Fig 6A). However, CtsB did not reduce cell death (Fig 6B and C). The results were the same even at higher (10 µM) Ca- 074 Me concentration (Fig 6D and E). This suggests that VLPs may induce the cytotoxicity by different mechanism next to lysosomal damage-associated cell death.

**Figure 6.**
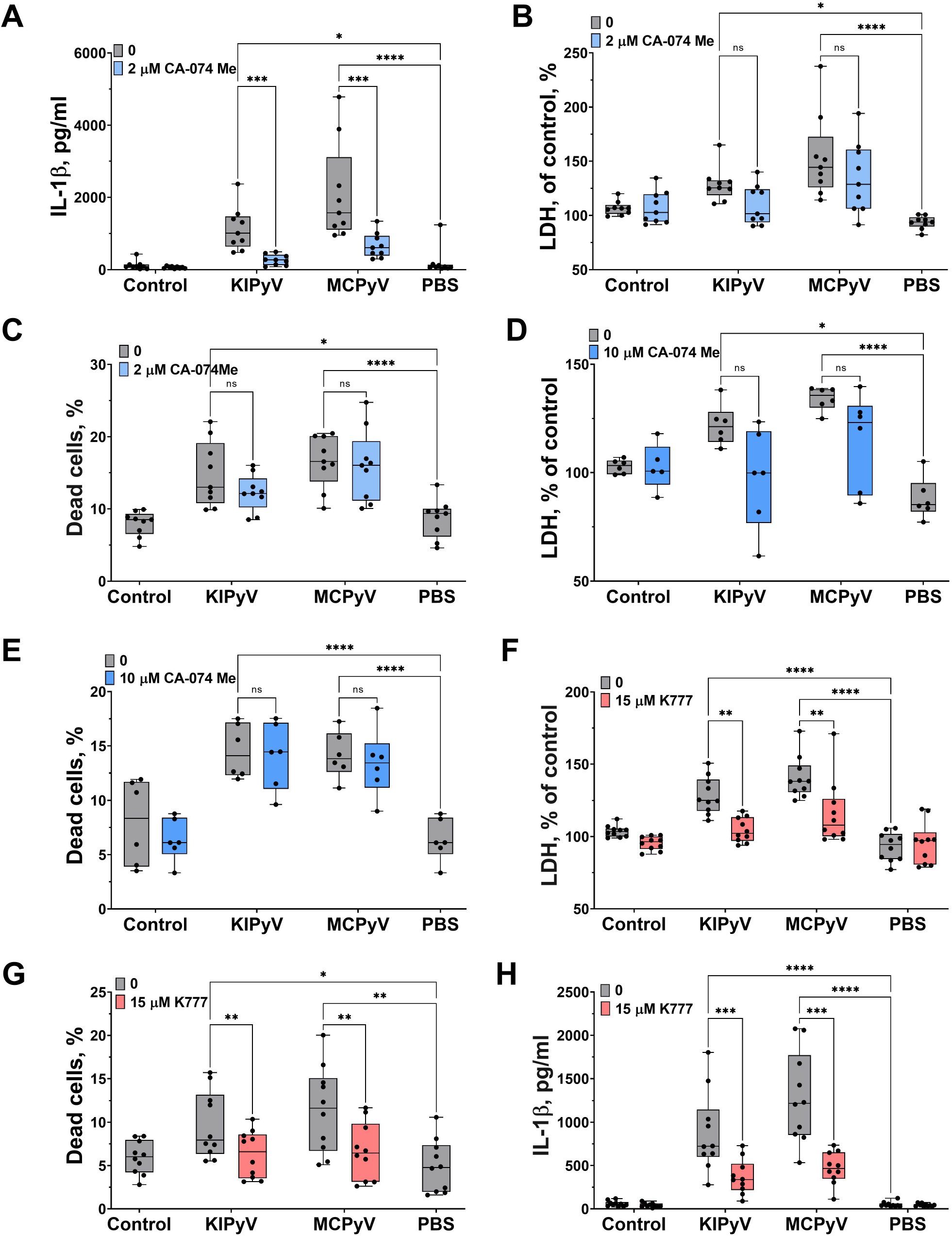
Cathepsin B inhibitor reduced VLP-induced IL-1β release while pan-cathepsin inhibitor (K777) decreased both IL-1β release and cell death in human THP-1 macrophages. A, H Macrophages were treated with PyV-derived VLPs (20 µg/ml) for 24 h. Inhibitors CA- 074 Me (2 or 10 μM) and K777 (15 μM) were added 30 min before treatment. (A, H) IL-1β secretion by ELISA. B-G Cytotoxicity assessed by (B, D, F) LDH assay and (C, E, G) PI and Hoechst nuclear staining. PI indicates dead cells (red) and Hoechst stains all cell nuclei (blue). Data information: Data are represented using box plots with dots showing independent experiments, *p < 0.05, **p < 0.01, ***p < 0.001, ****p < 0.0001, one-way ANOVA followed by Tukey’s multiple comparison test.

Then, we treated THP-1 cells with the pan-cathepsins inhibitor K777. We observed a significant decrease in cell death (Fig 6F-H). Staining of cell nuclei with PI to count dead cells confirmed that K777 inhibitor protected from cell death. Next, we measured IL-1β release and found that K777 significantly suppressed IL-1β secretion, however, not to control baseline. This indicates that K777 is a partial inhibitor of inflammasome activation induced by PyV-derived VLPs.

### PyV-derived VLPs induced inflammasome activation via K^+^ efflux

Lysosomal damage can induce further processes which consolidate inflammasome activation by VLPs as it was shown with inhaled particles, like silica and polystyrene nanoparticles (Rabolli *et al*., 2016a). The VLPs might also trigger different cell activation pathways leading to inflammasome activation. To prove this assumption, we investigated an alternative inflammasome activation pathway using K^+^ efflux inhibitor, glybenclamide. This inhibitor significantly blocked cell death according to LDH release data (Fig 7A). We also observed a significant decrease in IL-1β release after glybenclamide pre-treatment (Fig 7B).

**Figure 7.**
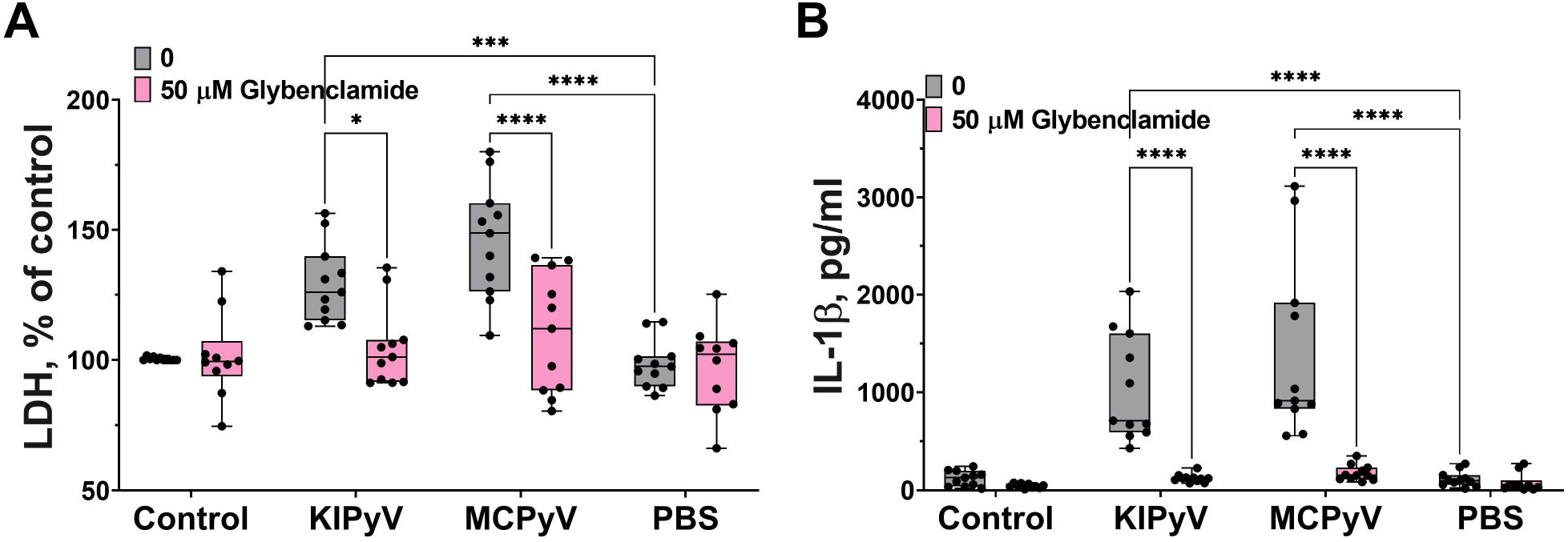
K^+^ ion efflux inhibitor (glybenclamide) reduced PyV VLP-induced cell death and IL-1β release in human THP-1 macrophages. A Macrophages were treated with PyV-derived VLPs (20 µg/ml) for 24 h. Inhibitor glybenclamide (50 μM) was added 30 min before treatment. (A) Cytotoxicity assessed by LDH assay. B IL-1β secretion determined by ELISA. Data information: Data are represented using box plots with dots showing independent experiments, *p < 0.05, **p < 0.01, ***p < 0.001, ****p < 0.0001, one-way ANOVA followed by Tukey’s multiple comparison test.

Taken together, these results reveal the complexity of inflammatory responses leading to inflammasome activation induced by PyV-derived VLPs in THP-1 macrophages. It is likely that several different mechanisms are involved in this process.

### Inflammasome activation by PyV-derived VLPs was confirmed in primary human macrophages

As cell lines may misrepresent real cell activation pattern, we used primary human macrophages derived from monocytes of peripheral blood mononuclear cells to confirm cell activation profile observed in THP-1 cells treated with PyV-derived VLPs. Activation of primary human macrophages with VLPs revealed a similar cell activation pattern to that observed in THP-1 macrophages – a significant increase in activated caspase-1 (Fig 8A and B), cell death (Fig 8C), and TNF-α and IL-1β release (Fig 8D and E). Moreover, NLRP3 inflammasome inhibitor MCC950 significantly reduced primary human macrophage activation and rescued from cell death (Fig 8A-C). As in experiments with THP-1 macrophages, not all primary human macrophages were activated by the VLPs and the percentage of dead cells did not exceed 20% (Figs 3E and 8C). According to FLICA assay indicating caspase-1 activation (Fig 8A and B), only a part of primary human macrophages was activated by VLPs. According to PI staining, primary macrophages having activated caspase-1 also were dead (Fig 8A) proving the pyroptotic cell death. These experiments indicate a similar pattern of PyV VLP-induced inflammation followed by inflammasome activation both in THP-1 cell line and in primary human macrophages.

**Figure 8.**
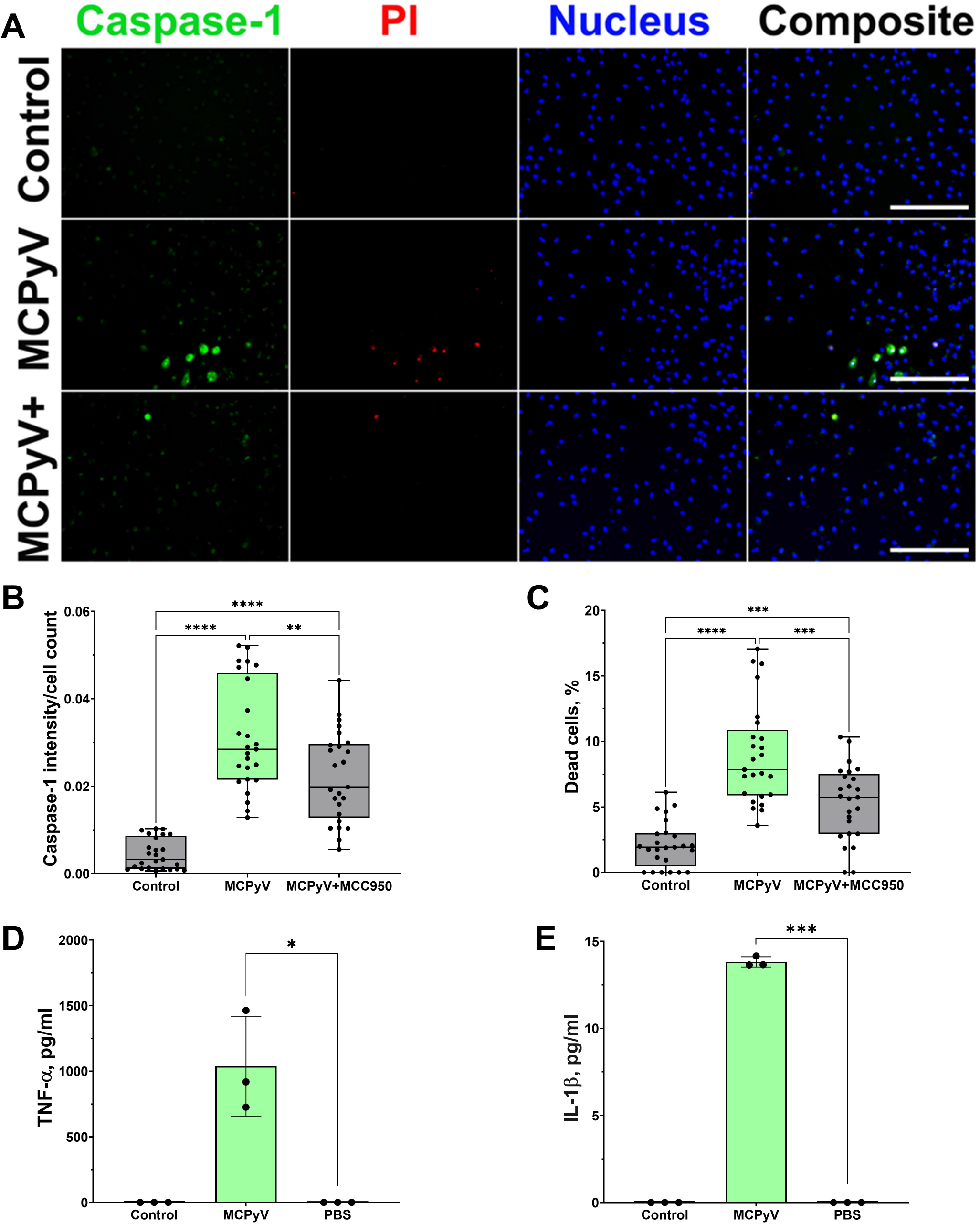
PyV-derived VLPs activated NLRP3 inflammasome in primary human macrophages-derived from peripheral blood mononuclear cells. A–C Macrophages were treated for 15 h with PyV-derived VLPs (20 µg/ml), inhibitor MCC950 (1 μM) was added 30 min before treatment. Cells were stained for activated caspase-1 using FLICA (green), dead cell nuclear stain PI (red) and nuclear stain Hoechst (blue) (A). The images were taken using 20× objective. Representative images of the staining are shown. The scale bars indicate 100 μm. (B) Quantification of dead cells, n = 25 photos per condition. (C) Caspase-1 quantification according to FLICA analysis, n = 25 photos per condition. D, E After 24 h treatment with PyV-derived VLPs collected supernatants were analysed by ELISA for (D) TNF-α and (E) IL-1β release, n = 3 technical repeats. Data information: N = 1 independent experiment. For (A–C) data were analysed using one-way ANOVA followed by Tukey’s multiple comparison test and represented using box plots with dots showing individual data points. For (D, E) Student’s t-test was used and data were represented using bar graphs (means±SD) with dots showing individual data points. *p < 0.05, **p < 0.01, ***p < 0.001, ****p < 0.0001.

We found that in some THP-1 cells the inflammasome is activated and in some is not based on data obtained from caspase-1 and ASC speck formation assays. To identify differences between these cells, we performed single-cell RNA sequencing (ScRNAseq) analysis. This analysis was also used to reveal whether KIPyV and MCPyV VLPs induce different cell activation states since we observed different cell activation levels according to inflammatory cytokine production.

### ScRNAseq reveals an overall conserved gene expression response to PyV-derived VLPs

*In vitro* differentiated THP-1 macrophages were subjected to stimulation by VLPs of KIPyV or MCPyV (and unstimulated control – PBS) for 15 h followed by single-cell RNAseq (Fig 9A). Upon removing transcriptomes with <900 total counts, 32,415 cells were retained, with >10,000 cells per condition. The mitochondrial gene count filter, conventionally used as a signature of dead cells (Luecken & Theis, 2019), was omitted to enable a comparison of the viability observed from scRNAseq data.

**Figure 9.**
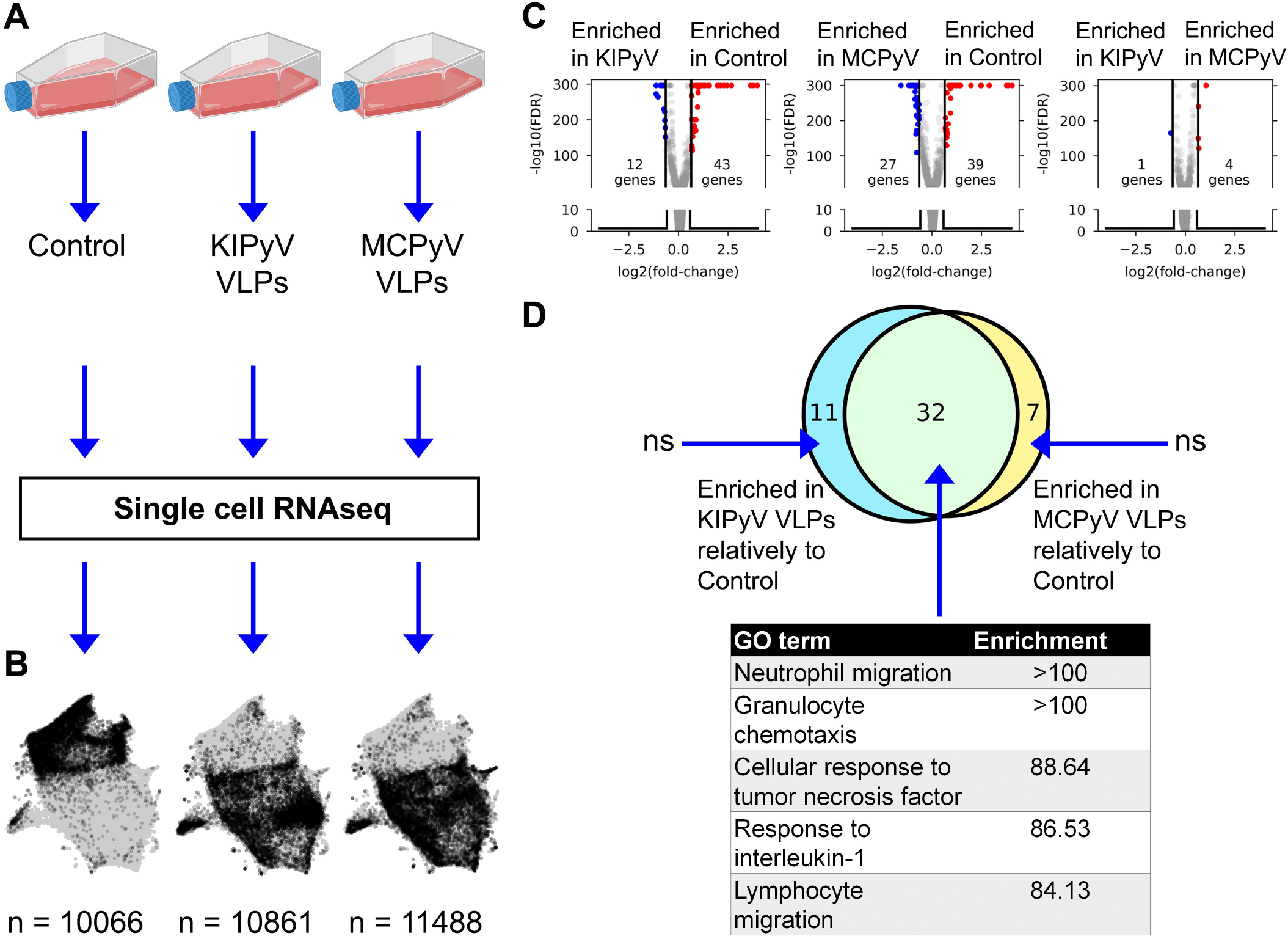
ScRNAseq gene expression profiling of THP-1 macrophages stimulated with PyV-derived VLPs. A Experiment outline. B UMAP representation of scRNAseq data. Grey dots denote cells from all conditions combined. Black dots highlight cells from a given condition. Numbers at the bottom indicate the number of single cell transcriptomes post-filtering. The conventional mitochondrial count filter (to remove dead cells) was intentionally omitted to enable the estimation of the fraction of dead cells. C Volcano plots showing bulk-like differential genes expression analysis results. Differentially expressed genes (DGEs) were defined as having an absolute fold-change > 1.5 and FDR < 0.05 (Mann-Whitney U test). D **Top:** Venn diagram showing the overlap between sets of genes enriched in KIPyV VLP-treated cells relatively to control, MCPyV VLP-treated cells relatively to control and MCPyV VLP-treated cells relatively to KIPyV VLPs. **Bottom**: GO enrichment analysis results for each subset of genes shown in the Venn diagram. N.s. – no significant GO term enrichment. Data information: To prepare (A) https://biorender.com/ was used.

Uniform Manifold Approximation and Projection (UMAP) visualization of scRNAseq data revealed an overall similar population structure between KIPyV and MCPyV VLP treated cells, and an apparent difference compared to the unstimulated control (Fig 9B). A bulk-like differential gene expression (DGE) analysis identified 43 upregulated and 12 downregulated genes in KIPyV VLP case vs the control (Fig 9C and Table EV1). The equivalent analysis of MCPyV VLP case vs control revealed 39 and 27 genes, respectively By contrast, a total of 5 differentially-expressed genes (DEGs) were identified when comparing MCPyV vs KIPyV VLP-treated cells. Most of the enriched genes compared to the control were the same regardless of the VLPs used for stimulation, and the 5 most-enriched terms in a GO gene set enrichment analysis of these common genes were associated with immune cell activation processes (migration, chemotaxis, response to TNF and IL-1) (Fig 9D and EV9, Table EV2).

### ScRNAseq reveals multiple subpopulations of THP-1 macrophages and changes in their abundance upon VLP stimulation

The UMAP visualization of scRNAseq data revealed a continuous structure with no clear boundaries between gene expression states. Upon interactive exploration (Weinreb *et al*, 2018) of gene expression patterns and Leiden clustering results at different resolution (i.e. different number of clusters), we chose to report on 9 populations (Fig 10A) of variable abundance across the 3 conditions (Fig 10B and C). For each cell population, we identified 20 most enriched genes (FDR<0.05, Mann-Whitney U test). Hierarchical clustering of the expression of these genes across all 9 populations revealed both distinct and overlapping gene expression signatures and the transcriptional relationship between populations (Fig 10D and E). The major split of the population dendrogram (Fig 10D) separates activated states (AI, AII, HA, IR) enriched after VLP stimulation (Fig 10F) from SS, RS, MA, and PA. The population of dead cells cluster with the latter but form a distant branch. Consistently with cell viability assays (Fig 3D and E), the fraction of dead cells, characterized by a high expression of mitochondrial genes (Fig 10E and Fig EV10A, Table EV3), increases with VLP stimulation from 2% to 5-6% (Fig 10C).

**Figure 10.**
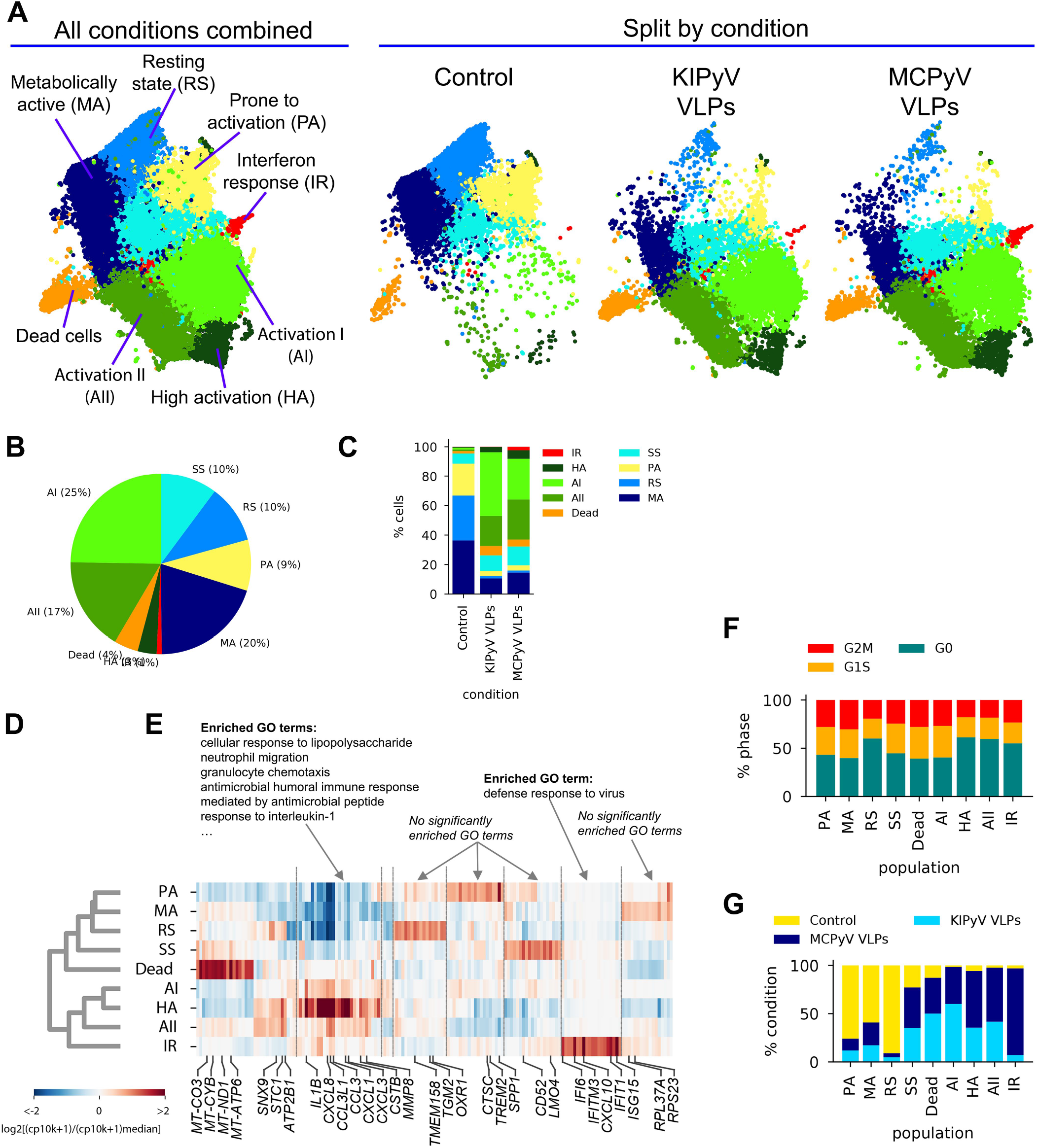
ScRNAseq revealed different cell populations within THP-1 macrophages treated with PyV-derived VLPs. A UMAP coloured by cell population annotation either combining all conditions (left) or split by condition (right). B, C Percentages of cells per population for all data combined (B) or split by condition (C). D, E Results of population-enriched gene identification, including a population dendrogram (D) based on hierarchical clustering (correlation distance measure, average linkage) by enriched gene expression (E). Selected genes are highlighted at the bottom. Results of GO enrichment analysis on selected genes groups are shown on top. CP10K – counts per 10,000. F Relative abundances of each population by condition. As this analysis is sensitive to the total number of cells sampled by condition, cell counts from each condition were first normalized to 10,000. G Relative abundance of cells at different cell cycle stages defined based on gene expression scoring.

None of the states was completely quiescent, i.e., with all cells in the G0 stage, as suggested by a classification into cell cycle stages based on gene expression, although different populations showed a variable fraction of cells in G1/M/G2/S stages (Fig 10G).

The cell cluster most uniformly represented across the three conditions (PBS, KIPyV and MCPyV VLPs) was called steady state (SS) (Figs 10A and 10F). Among its enriched genes are *CD52*, *LMO4*, *CHI3L1*, *COX5B* (Figs. 10E and EV10Bb, Table EV3). The function of most of proteins encoded by these genes is unclear. CD52 is a glycoprotein of an unknown function. A recent study showed the ability of soluble CD52 to suppress inflammatory cytokine production by inhibiting TLR-induced NF-κB activation in macrophages, monocytes, and dendritic cells (Rashidi *et al*, 2018). *COX5B* encodes 5B subunit of Cytochrome C Oxidase, an essential mitochondrial respiratory chain enzyme (Galati *et al*, 2009). An increased expression of *COX5B* may be related to an increased cellular respiration. *LMO4*-encoded protein may play a role as an oncogene targeting TGF-β signalling pathway (Lu *et al*, 2006). Overall, the phenotype of SS population is unclear as the functions of most enriched genes in macrophages are unknown. In addition, GO enrichment analysis did not show any relation to known GO terms (Fig 10E). Therefore, this population could be described as a new possible phenotype of THP-1 cell line.

One cell cluster, the relative abundance of which changed upon cell activation, was called resting state (RS) (Fig 10A). This cell population did not express inflammatory molecules and the enriched genes were related to usual cellular processes. Even so, this cluster distinguished oneself by high enrichment of *ARHGAP18*, *TMEM158*, *TGM2*, *OXR*, and *FN1*, which have vital or still unknown functions (Figs 10E and EV10C, Table EV3). ARHGAP18 encodes Rho GTPase Activating Protein 18, which is essential for actin remodelling, thus, it is important for cell migration and controls cell shape (Maeda *et al*, 2011). *TMEM158* encodes Transmembrane Protein 158 which biological function is unclear, although its involvement in activation of Ras pathway was shown. It was reported that TMEM158 enhanced proliferation and migration of cancer cells (Cheng *et al*, 2015; Liu *et al*, 2020). For example, another transmembrane protein TMEM119 was identified as a marker of brain-resident macrophage, called microglia, marker, representing microglia with homeostatic properties (Butovsky *et al*, 2014; Satoh *et al*, 2016). Noticeably, *TMEM158* expression decreased after activation with VLPs addressing its presence in ramified macrophages (Fig EV10I). *TGM2* encodes Transglutaminase 2 implicated in cell death pathways (Mastroberardino *et al*, 2002). *OXR1* encodes Oxidation Resistance Protein 1, which is involved in protection from oxidative stress and is important for lysosomal function (Volkert *et al*, 2000; Wang *et al*, 2019). *FN1* encodes fibronectin 1 known for its function in cell adhesion, cell motility, and maintenance of cell shape (Hynes, 1986; Kornblihtt & Gutman, 1988). Therefore, expression of these genes indicates restful cells.

Another cell cluster that changed upon cell activation was called metabolically active (MA) (Fig 10A). It had a higher expression of mitochondrial and ribosomal genes, for example, *MT-CO3*, *MT-CYB*, *RPL37A*, and *RPS23* (Figs. 10E and EV10d). The MA cluster did not show expression of genes related to inflammatory cell activation, but the observed gene expression profile indicates increased cellular respiration that is related to high cell metabolic activity (Osellame *et al*, 2012). Enrichment in ribosomal genes characterized protein production indicating metabolic cell activity.

In the control condition (PBS) we found a cluster in which the cells seemed to be activated *a priori*. It was named prone to activation (PA) (Fig 10A). Higher expression of *SPP1*, *CTSC*, *TREM2*, *S100A4*, and *IL1B* was detected in some cells of this cluster (Figs 10E, EV10E and EV10F, Table EV3). It is possible that these cells showed a delayed response to phorbol 12-myristate 13-acetate (PMA) used for THP-1 differentiation. This cell cluster disappeared after treatment with VLPs. *CTSC* encodes lysosomal protease Cathepsin C. As other cathepsins, CTSC is important for cargo degradation (Kominami *et al*, 1992). CTSC was also shown to be necessary for activating serine proteases since its absence altered extracellular IL-1β activation mediated by serine proteases (Rabolli *et al*, 2016b). In addition, upregulation of CTSC mediates macrophage polarization to inflammatory phenotype (Alam *et al*, 2019; Liu *et al*, 2019). *S100A4* encodes calcium-binding protein S100A4 involved in inflammatory reactions (Hansen *et al*, 2015; Helfman *et al*, 2005). *SPP1* encodes Secreted Phosphoprotein 1 that is chemotactic, induces IFN-γ and IL-12 production, and promotes cell survival (Wang & Denhardt, 2008). *TREM2* encodes Triggering Receptor Expressed on Myeloid Cells 2 that triggers secretion of inflammatory molecules (Bouchon *et al*, 2001; Kobayashi *et al*, 2016), although, anti-inflammatory effect of TREM2 was demonstrated in macrophages lacking TREM2 as toll-like receptor stimulation induced higher pro-inflammatory cytokine secretion (Turnbull *et al*, 2006). Soluble TREM2 was identified as an activator of inflammatory response (Zhong *et al*, 2017). It enhances phagocytosis as its loss impairs cellular uptake of various substrates, such as cellular debris (Takahashi *et al*, 2005), bacteria (N’Diaye *et al*, 2009) or amyloid-beta aggregates (Jiang *et al*, 2014). Overall, PA cells are differently activated than other THP-1 cell populations (AI, AII, HA, IF) and they distinguish themselves as having inflammation-related and phagocytic cell properties.

The cluster of cells formed upon VLP stimulation was defined as a high activation (HA) state (Fig 10A). This cluster was more abundant in MCPyV VLP-treated cells than in KIPyV VLP-treated cells (Fig 10F). This cell population was characterized with the highest expression of *IL1B* and chemokine genes, such as *CXCL1*, *CXCL3*, *CXCL8*, *CCL3*, *CCL20*, *CCL4L2*, and *CSTB* (Figs. 10E and EV10F, Table EV3). It is known that IL1B is expressed at extremely high levels in myeloid-derived cells in response to microbial invasion and tissue injury (Adamik *et al*, 2013). The product of *IL1B* is pro-inflammatory cytokine IL-1β, a key mediator of inflammation and one of the main indicators of NLRP3 inflammasome assembly (Tominaga *et al*, 2000). IL-1β induces the production of chemokines and proteases, such as matrix metalloproteinase, to attract other immune cells to the infection site. Furthermore, secretion of chemokines is reduced in NLRP3-deficient mice demonstrating the importance of inflammasome activation in chemotaxis (Shimizu *et al*, 2019). IL-1β is a product of inflammasome activation, thus, factors stimulating the inflammasome also recruit immune cells. *CXCL1*, *CXCL3*, and *CXCL8* encode members of the CXC subfamily of chemokines, the chemoattractants for neutrophils (Raman *et al*, 2011). CXCL8 also known as IL-8 has no homologs in rats or mice and is a significant component of inflammation-mediated processes as it attracts neutrophils, basophils, and T-cells to the site of infection and promotes endothelial cell migration, invasion, and proliferation (Schutyser *et al*, 2002). HA state is also rich in genes of other chemoattractants CCL3 and CCL20. CCL3, also known as macrophage inflammatory protein 1 alpha, is induced by NF-κB signalling pathway and triggers inflammatory reactions (Cook, 1996). CCL20 acts as a ligand for C-C chemokine receptor CCR6 that induces a strong chemotactic response and plays an important role at skin and mucosal surfaces under homeostatic and inflammatory conditions (Ito *et al*, 2011). HA cluster was also highly enriched in *CSTB* which encodes cystatin B, a cysteine protease inhibitor known to interact with cathepsin B (Pavlova *et al*, 2000) and considered to protect from cathepsin leakage from damaged lysosomes (Mrschtik & Ryan, 2015). *CSTB* expression reveals possible lysosomal damage induced by VLPs. Overall, highly enriched genes of HA state showed a strong inflammatory response in VLP-treated macrophages.

Other cell clusters formed upon cell activation were named the Activation I (AI) and Activation II (AII) states (Fig 10A). AI cluster was more characteristic for KIPyV than MCPyV VLP-treated cells and AII state *vice versa* (Fig 10F). Both clusters had enriched inflammation-related genes similarly to HA state but at lower levels (Fig 10E). AI state was characterized by high *IL1B*, *CXCL8*, *CCL3L1*, and *CCL3* expression (Figs. 10E and EV10G, Table EV3). AII had relatively low expression of *IL1B* but the enrichment in *CXCL8*, *CCL3L1* and *CCL3* was similar to AI. In general, the same genes were enriched in HA, AI, and AII states, although the expression levels were different. Furthermore, HA and AII were different from AI state as HA and AII had enrichment in *STC1*, *MMP8*, *ATP2B1*, *FTH1*, and *SNX9* genes. Interestingly, enrichment in some of these genes, like *ATP2B1* and SNX9, was similar to RS cluster. *STC1* is known to encode Stanniocalcin 1, a secreted glycoprotein involved in inflammation and carcinogenesis (Chang *et al*, 2003) that possibly serves as a negative mediator of inflammation (Leung & Wong, 2021). Interestingly, HA state, which was highly enriched in pro-inflammatory cytokines, also expressed high levels of *STC1*. MMP8 is a Matrix Metalloproteinase-8 that cleaves collagen, some cell adhesion proteins, growth factors and chemokines (Van Lint & Libert, 2006), and promotes polarization of macrophages into alternatively activated (M2) macrophages (Wen *et al*, 2015). Therefore, AII and HA states describe a group of activated macrophages expressing inflammation-related genes and a couple of inflammation suppressors. *SNX9* encodes Sorting Nexin 9 involved is in intracellular trafficking (Kurten *et al*, 1996), and regulate clathrin-mediated endocytosis (Carlton *et al*, 2005). It also plays a role in inflammatory reactions (Ish-Shalom *et al*, 2016) and regulation of micropinocytosis – endocytosis pathway (Wang *et al*, 2010). In innate immune cells this endocytosis pathway may function for the delivery of antigens to their respective intracellular pattern recognition receptors (Canton, 2018). Since endocytosis pathway of polyomavirus VLPs is unclear, SNX9 may contribute to intracellular recognition of VLPs. *ATP2B1* encodes Plasma Membrane Calcium ATPase 1 (PMCA1) important in maintaining cytosolic Ca^2+^ for physiological cell functions (Brini & Carafoli, 2011). Ca2+ mobilization is critical for NLRP3 inflammasome activation (Murakami *et al*, 2012). Thus, calcium pumps could be implicated in inflammasome signalling detected after VLP treatment. *FTH1* encodes the heavy subunit of ferritin which is essential to store iron inside cells (Gozzelino & Soares, 2014). Macrophages are central players in iron metabolism as they recycle senescent erythrocytes and modulate iron availability as part of host protective mechanisms (Soares & Hamza, 2016). FTH1 is crucial in protection against iron-induced oxidative stress and cell death as it is involved in decreasing peroxide formation from iron (Mesquita *et al*, 2020). In summary, HA and AII clusters next to inflammation-related genes also expressed higher levels of genes encoding ion channels and proteins involved in endocytosis contrary to AI state.

### While most population and gene expression changes are conserved, interferon-response population is highly enriched upon activation with MCPyV-derived VLPs

Analysis of the previously identified (Fig 9C and D) global gene expression changes at the individual population level revealed very similar gene expression patterns between KIPyV and MCPyV VLP-treated THP-1 cells (Figs 11A-C and EV11A). A population abundance fold-change analysis emphasized the overall similar phenotype of the two VLP-treatment conditions (Fig 11D), while abundance differences in abundance relatively to control exceeded 30x (Fig 11E, F). However, a notable exception was the interferon-response population (IR), highly enriched after MCPyV VLP stimulation compared to KIPyV VLP (Fig 11D), and characterized by the expression of *CXCL10*, *ISG15*, *IFITIM3*, *IFIT1-3*, *IFI6*, *CXCL11* and other genes (Figs 10E and EV10H, Table EV3), known to be involved in inhibition of viral invasion. Interestingly, we found an increase in interferon-inducible gene expression only after treatment with MCPyV VLPs. We assume that macrophage engagement with viral antigens itself mediates anti-viral activity in macrophages without external stimulation, such as lipopolysaccharide. In addition, the expression of IR-specific genes was essentially absent in control cells and after treatment with KIPyV VLPs (Fig EV10B). This suggests differences in pathogen-associated molecular patterns derived from human PyVs.

**Figure 11.**
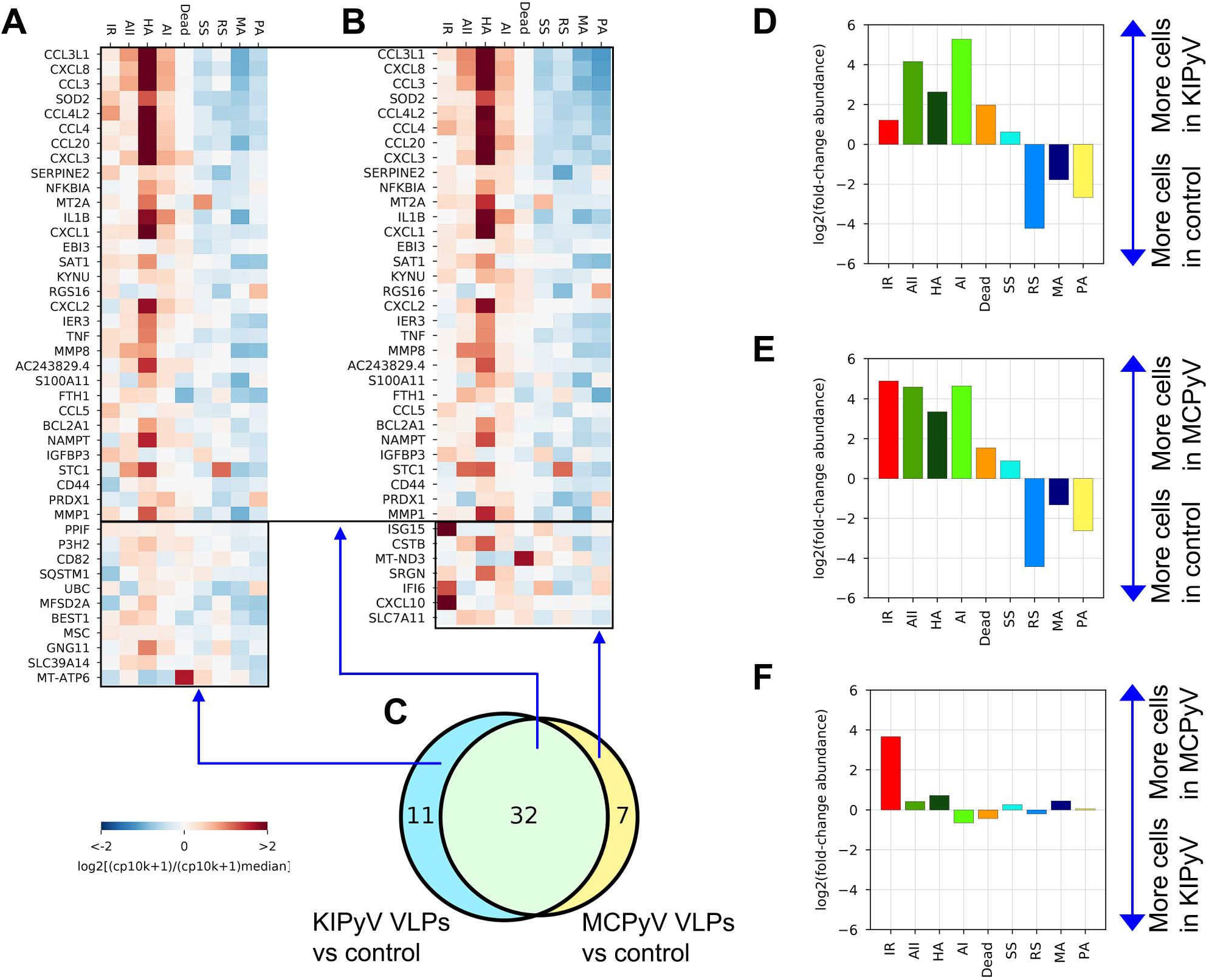
Genes enriched in both KIPyV and MCPyV VLP-treated cells have a very similar expression pattern across the populations observed, except for the interferon-response (IR) population. A Expression of genes identified as enriched in KIPyV VLP-treated macrophages in the bulk-like analysis (Fig 9C and D) at the individual population level in KIPyV VLP-treated macrophages. B The equivalent of A for the MCPyV VLP-treatment condition. C Same Venn diagram as in Fig 9D. D-F Fold-changes in population abundance in pairwise comparisons of the three conditions (Control, KIPyV VLP, MCPyV VPL).

In detail, most of IR cluster genes are induced by interferon (IFN) and related to cell response during viral infection. *IFITMs* encode Interferon-Induced Transmembrane Proteins, that limit viral infection via entry pathway (Zhao *et al*, 2018). In interferon-stimulated cells IFITM3 is localized on endocytic vesicles which fuse with incoming virus particles and enhance their trafficking to lysosomes (Spence *et al*, 2019). Recent study revealed that IFITM3 is able to inhibit SARS-CoV-2 infection *in vitro* (Shi *et al*, 2021). *IFITs* encode Interferon-Induced Proteins with Tetratricopeptide repeats that block viral infection by interacting with factors responsible for virus replication (Zhou *et al*, 2013). *IFI6* encodes Interferon-α Inducible Protein 6. The exact function of this protein is still unclear. The family of IFI6 genes encodes mitochondrial proteins implicated in inhibition of apoptosis (Cheriyath *et al*, 2011). IFI6 was also shown to inhibit DNA replication in hepatitis B virus (Sajid *et al*, 2021). *ISG15* (Interferon Stimulated Gene 15) encodes ubiquitin-like protein that is induced by type I interferons and involved in various processes related to host antiviral response, acting as extracellular and intracellular signalling protein (Perng & Lenschow, 2018). Its upregulation was detected in macrophages after infection by vaccinia and SARS-CoV-2 viruses (Sanyal, 2020; Yángüez *et al*, 2013). This protein is likely to be an important factor implicated in cytokine storms triggered by SARS-CoV-2. ISG15 deletion in bone marrow-derived macrophages induced mitochondrial dysfunction and altered nitric oxide production exposing its importance in regulating mitochondrial function (Baldanta *et al*, 2017). Other genes rich in IR state encode CXCL10 and CXCL11, but the latter expression was detected only in several cells. *CXCL10* is known as early interferon response gene encoding CXCL10 with inflammation-related pleiotropic effects (Booth *et al*, 2002). CXCL11 participates in inflammatory reactions and acts as a chemoattractant of activated T cells (Gasperini *et al*, 1999). CXCL10 and CXCL11 are thought to be key mediators of the cytokine storm in immune response to SARS-CoV-2 infection (Callahan *et al*, 2021). Overall, high expression of IR-specific genes indicates cellular response to interferon induced in THP-1 cells by VLP treatment.

## Discussion

In our study we focused on the inflammatory response of human macrophages treated with viral oligomeric proteins. Cellular response to nanoparticles has been explored previously covering different mechanisms. It was demonstrated that polymeric nanoparticles induced NLRP3 inflammasome activation dependent on cathepsin B in macrophages (Vaine *et al*, 2013). Another study showed that oligomeric proteins, Aβ fibrils, cause lysosomal damage and induce inflammasome activation (Halle *et al*, 2008). Our previous study showed that Aβ oligomers and protofibrils activated the NLRP3 inflammasome in microglia cells (Luciunaite *et al*.). In addition, tau oligomers and α-synuclein fibrils were shown to trigger an inflammatory response in microglia (Ising *et al*., 2019; Pike *et al*, 2021). The latter reports demonstrated inflammasome activation in macrophages by endogenous pathogenic protein oligomers. However, there are limited data on molecular mechanisms how oligomeric proteins activate NLRP3 inflammasome. To address this question, we investigated the ability of exogenous oligomeric proteins of viral origin to induce inflammatory responses in human macrophages.

In our study, we have used different viral oligomeric proteins – filamentous NLPs of measles and mumps viruses and spherical VLPs of KIPyV and MCPyV. We did not observe any inflammatory response to NLPs, although, NLRP3 inflammasome activation was demonstrated to be induced by N protein of SARS-CoV-2 (Pan *et al*, 2021). Interestingly, N protein of SARS-CoV-2 inhibits the cleavage of gasdermine D, which forms pores in the membrane, reducing IL-1β secretion and pyroptosis (Ma *et al*, 2021). In our study, only PyV-derived VLPs induced inflammatory response of human macrophages followed by NLRP3 inflammasome activation that was demonstrated by TNF-α and IL-1β release, cytotoxicity induction, caspase-1 activation and ASC speck formation. Specific NLRP3 inflammasome inhibitor MCC950 blocked the detected activation signal proving PyV-derived VLPs as a trigger of NLRP3 inflammasome. In addition, VLPs induced secretion of other inflammatory cytokines TNF-α and IL-6 suggesting the engagement of NF-κB signalling pathway in the activated macrophages.

NLRP3 inflammasome can be activated by different mechanisms, such as changes in intracellular ion concentration, mitochondrial or lysosomal damage followed by cathepsins release (Kelley *et al*, 2019). First, we studied inflammasome activation mechanism related to lysosomal damage since PyV-derived VLPs are phagocytosed particles. The VLP-induced inflammasome activation signal was significantly reduced by cathepsin B and pan-cathepsin inhibitors. However, the inhibitory effect was not complete compared to control. Cathepsin B inhibitor reduced only IL-1β release and had no effect on cell death. In the case of MCPyV-derived VLPs, IL-1β secretion and cell death did not drop to control baseline even using pan-cathepsin inhibitor. In the case of KIPyV-derived VLPs, pan-cathepsin inhibitor completely reduced cell death, contrary to IL-1β release. This suggests that VLP size and possibly other structural features of VLPs may define cell activation profile. Viral capsids of KIPyV and MCPyV are structurally different, thus, they may interact with different cellular receptors. In addition, the hemagglutination ability was demonstrated only for MCPyV indicating different glycoproteins on KIPyV and MCPyV capsid surface (Neu *et al*, 2012; Neu *et al*, 2011).

There are controversial data on particle-induced inflammasome activation. Some of them show that nanoparticles induce NLRP3 inflammasome activation via phagosomal destabilisation (Hornung *et al*, 2008). Other studies reveal that different nanoparticles induce different mechanisms depending on the composition and structure of the particles (Rashidi *et al*, 2020). For example, cholesterol crystals activate NLRP3 inflammasome, however, inhibitors of cathepsins reduce only IL-1β release and do not change the level of cell death. It is assumed that inflammasome can be activated by further inflammation mediators, such as reactive oxygen species and K^+^ ion efflux. On the other hand, cell death due to cell membrane damage could be an irreversible process. In addition, the role of cathepsins in inflammasome activation is not fully understood. For example, it was recently shown that cathepsins can induce activation of pore-forming protein gasdermin D (Selkrig *et al*, 2020).

We assumed that PyV-derived VLPs induced inflammasome activation via lysosomal damage and investigated the underlying mechanisms related to VLP-induced cell death, in particular the mechanism of K^+^ ion efflux. We demonstrated that a specific inhibitor of K^+^ ion efflux significantly reduced cell death and IL-1β release suggesting the complexity of macrophage activation by PyV-derived VLPs. In addition, studies on the possible outcome of lysosomal leakage indicate that effector molecules such as cathepsins released after the permeabilisation of lysosomal membrane may activate the inflammasome by several mechanisms, including K^+^ ion efflux and intracellular Ca^2+^ initiated signalling events (Stahl- Meyer *et al*, 2021). Cathepsins can also induce synthesis of IL-1β precursor leading to a higher release of inflammatory cytokines and involvement of cathepsins in NF-κB signalling (Orlowski *et al*, 2015). This demonstrates that inflammasome activation by phagocytosed particles is beyond the classical mechanism.

Finally, we proved the inflammatory response observed in human macrophage cell line THP-1 to PyV-derived VLPs using primary human macrophages. The same cellular events were identified both in THP-1 cell line and the primary human macrophages. Activated caspase-1 was detected in dead primary macrophages suggesting the pyroptotic cell death. As in THP-1 experiments, only a part of the primary macrophages was activated by the VLPs. To conclude, the inflammatory response of cells treated with PyV-derived VLPs demonstrates the capability of large-sized multimeric protein particles to induce a similar cell activation pattern both in macrophage cell line and primary human macrophages.

From *in vitro* studies it is well known that strong inducers of inflammation, such as lipopolysaccharides or nigericin, can induce high cell activation states. However, other cell activators usually mediate modest cellular response. For example, in previous research we showed lower inflammasome activation by Aβ oligomers compared to classical NLRP3 inflammasome inducer nigericin (Luciunaite *et al*.). However, when Aβ-activated cells were observed microscopically it was clear that not all of them exhibited similar level of inflammasome activation. It raises a question on the activation state of separate cells. The presence of heterogeneous cell populations was demonstrated several times by other groups. For example, brain macrophages, microglia, are at different cell activation states depending on the distance from amyloid-beta (Aβ) plaque in the brain of Alzheimer’s disease patients (Swanson *et al*, 2020). Microglia with different phagocytic capabilities were also identified in Alzheimer’s disease-relevant mouse model. Interestingly, microglia non-containing Aβ had more changes in expression of genes associated with accelerated ageing process than microglia with Aβ content (Grubman *et al*, 2021). In addition, microglia are heterogeneous in healthy brain and cells enriched in inflammation-related genes are present throughout the lifespan and rises up in the aged brain (Hammond *et al*, 2019). ScRNAseq analysis of peripheral blood mononuclear cell culture revealed that monocytes respond differently to lipopolysaccharide (Lawlor *et al*, 2021). One cell population expressed pro-inflammatory alarmins and chemokines to attract immune cells while another was enriched in later inflammation-related and anti-inflammatory genes to either enhance or terminate inflammatory reaction. In our study, IL-1β and TNF-α release data showed overall cell activation by PyV-derived VLPs, however, caspase-1, ASC speck and cell viability assays allowed the identification of differently activated cells. Therefore, we performed a single-cell analysis to identify gene expression differences in cells which primarily were expected to be homogenous – THP-1 cell culture model.

ScRNAseq analysis showed highly divergent cell activation pattern after treatment with PyV-derived VLPs. Four activated THP-1 cell populations varying in expression of inflammation-related genes, among them IL1B and chemokine genes, were identified. IL-1 family cytokines are known to induce the production of chemokines (Dinarello, 2018). The stress response and prolonged inflammation also lead to recruitment of immune cells (Carta *et al*, 2017). Therefore, increased chemokine expression in cells, enriched in IL-1β and other inflammation mediators, indicates a strong inflammatory response. After VLP treatment, one cell cluster was highly enriched in inflammation-related genes (HA state), including IL-1β. This population was also enriched in CSTB which encodes cystatin B, an inhibitor of cathepsin B. It possibly copes with cathepsins released after lysosomal leakage (Mrschtik & Ryan, 2015). Therefore, CSTB expression verifies lysosomal damage induced by VLPs. This cell cluster may represent a population of inflammasome-activated cells, which we revealed according to ASC speck formation, cell viability and caspase-1 assays.

Interestingly, two populations of activated cells, including the HA cell cluster, also expressed negative mediators of inflammation next to inflammation-related genes. This may indicate a prolonged cell activation state when self-protecting genes were switched on to avoid hyperinflammation. It is known that after inflammatory response follows the resolution phase when anti-inflammatory molecules are secreted (Schett & Neurath, 2018).

Comparing KIPyV and MCPyV VLP-induced cellular response, we found the expression of interferon response-related genes only after MCPyV VLP treatment. This extends known KIPyV and MCPyV differences. Among interferon-response genes were CXCL10 and IGS15, involved in host antiviral response and found to induce cytokine storm in SARS-CoV-2 infected patients (Callahan *et al*., 2021; Sanyal, 2020). Interferon-induced proteins participate in inflammatory reactions and attract different cells of innate and adaptive immunity. Interferon-stimulated genes can be triggered not only by viral antigens directly but also by cellular stress response, thus, their encoded proteins may have a broader biological function (Fensterl & Sen, 2015).

Summarizing, single cell analysis revealed the presence of heterogeneous cell populations in the *in vitro* cell culture model, THP-1 macrophages, which might be expected to be a homogenous cell culture. Even more complex cell activation patterns might be predicted *in vivo*. Although the molecular mechanisms behind different cell activation states are unknown, our study shows that some cells respond to the activating agent by inflammatory reactions while other cells remain unaffected. Assuming the inflammasome activation by protein oligomers as a result of phagocytosis-related process, we suppose that some cells are capable to degrade their cargo and some not, which may lead to lysosomal disruption.

Inactivated viruses and recombinant viral proteins are broadly used for vaccination (Pollard & Bijker, 2020). However, the mechanisms of the immune response and especially activation of the innate immunity components induced by viral proteins are barely investigated. Therefore, the results of our study on PyV-derived VLPs as potent inducers of the inflammatory response in macrophages as compared to recombinant filamentous NLPs that are incapable to trigger the inflammation would broaden the understanding of the interaction of viral proteins with innate immune cells. Our study suggests that not all viral proteins can trigger the inflammatory response and their structural properties are one of the factors defining cell activation pathway.

## Materials and Methods

### Materials

Dulbecco’s modified Eagle’s medium (DMEM; cat# 31966047), Roswell Park Memorial Institute 1640 medium (RPMI, cat#61870044), FluoroBrite DMEM (cat#A1896701), fetal bovine serum (FBS; cat# A3840402), penicillin/streptomycin (P/S; cat#15140122), Dulbecco’s Phosphate Buffered Saline (PBS; cat#14190250), cell dissociation reagent TrypLE™ Express Enzyme (cat#12604021) were obtained from Gibco, ThermoFischer Scientific. Cell culture plates: T75 culture flasks Cell Culture Treated EasYFlasks (cat#156499) were from Nunc, ThermoFischer Scientific; TPP Multi-well tissue culture plates (cat# 92012, cat#92024, cat#92048) were from TPP Techno Plastic Products AG; IbiTreat 96-well μ-plates (cat#89626) were from Ibidi. LPS (cat#tlrl-eblps), nigericin (cat#tlrnig), MCC950 (cat#inh-mcc), normocin (cat#ant-nr-1) and zeocin (cat#ant-zn-05) were from InvivoGen. K777 [K11777] (cat#AG-CR1-0158-M001) was from Adipogen. CA-074 Me (cat#A8239) was from ApexBio Technology. LDH cytotoxicity detection kit (cat#11644793001) was from Roche Diagnostics, Sigma-Aldrich by Merck. Phorbol 12- myristate 13-acetate (PMA, cat# P1585-1MG) was obtained from Sigma-Aldrich by Merck. Propidium Iodide (PI; cat#638), Hoechst33342 (Hoechst, cat#639) and FAM-FLICA® Caspase-1 Assay Kit (containing FLICA reagent FAM-YVAD-FMK – caspase-1 inhibitor probe; cat#98) were obtained from ImmunoChemistry Technologies. Dimethylsulfoxide (DMSO; cat#A3672) was from PanReac AppliChem and the ITW Reagents. Human IL-1 beta Uncoated ELISA Kit (cat# 88-7261-77) and TNF alpha Uncoated ELISA Kit (cat#88- 7346-86), IL-6 Uncoated ELISA Kit (cat#88-7066), IL-10 Uncoated ELISA Kit (cat#88- 7106), Phosphate-Buffered Saline (10X) pH 7.4 (cat#AM9624), UltraPure DNase/RNase-Free Distilled Water (cat#10977035) were from Invitrogen, ThermoFischer Scientific. Tween-20 (cat# 9127.1) and sulphuric acid (H_2_SO_4_, cat#X873.1) were from CarlRoth. Chemiluminescent substrate – SuperSignal West Femto Maximum Sensitivity Substrate (cat#34094) was from ThermoFisher Scientific. NextSeq 500/550 High Output Kit v2.5 (cat#20024906) was obtained from Illumina.

### Cell lines

Human cell line THP-1 was kindly provided by prof. Linas Mažutis (Vilnius University, Vilnius, Lithuania). Cells were propagated in RPMI 1640 + 10% FBS + 100 U/mL of P/S and were split twice a week by ratio 1:5 to 1:10.

Human cell line THP-1 monocytes – ASC speck reporter cells were purchased from Invivogen (#thp-ascgfp, Invivogen, France) and called THP-1-ASC-GFP. These cells express ASC protein fused to GFP. Cells were propagated in RPMI 1640 + 10% FBS + 100 U/mL of P/S + 100 µg/ml of zeocin + 100 µg/ml normocin and were split twice a week by ratio 1:5 to 1:10. During cell differentiation to macrophages and treatment zeocin and normocin were not used.

### Human macrophage cell culture

Human macrophage cell cultures were prepared by differentiation of THP-1 cells (Chanput *et al*, 2014). The cells were seeded in the 24 well plate at a density of 0.125x10^6^/well using RPMI 1640 medium supplemented with 10% FBS and 1% P/S and differentiated to macrophages using 100 ng/mL of PMA. After 48 h of differentiation the medium was replaced with the fresh medium without PMA and cells were left to rest for another 24 h. After the rest period the cells are differentiated into macrophages and were used in experiments with viral proteins. These macrophage-like cells were used in the study and called THP-1 macrophages.

THP-1 macrophages were washed once with serum-free RPMI and treated with viral proteins for 24 h. Viral proteins were prepared in PBS, so, control when PBS was added instead of viral proteins was used. As a positive control, the inflammasome inducer nigericin was used at 10 μM concentration. MCC950, which selectively inhibits the NLRP3 inflammasome, was used at 1 μM concentration and added 30 min before the treatment. Inhibitors of cathepsins were used with the following concentrations: CA-074 Me at 2 μM and 10 μM, K777 at 15 μM, and added 30 min before the treatment. Another inhibitor, glybenclamide, which blocks K^+^ efflux, was used at 50 µM concentration, 30 min before the treatment. After incubation, cell culture supernatants were collected and stored at -20 °C for further cytokine analysis. Supernatants for LDH assay were used instantly.

Primary human macrophages were purchased from Lonza (#4W-700). Human macrophages were derived from CD14^+^ human monocytes of one donor. Cryopreserved cells were thawed and cultured for two days in RPMI 1640 medium supplemented with 10% FBS and 1% P/S before treatment. The cells were treated as THP-1 macrophages in serum-free RPMI. Cells for FLICA assay were plated in Ibidi 96-well μ-plate and for ELISA in TPP 24-well plate.

### Viral proteins

Macrophages were activated with recombinant viral proteins (20 µg/ml) representing various oligomeric shapes and forms. KI polyomavirus recombinant major capsid protein VP1 (KIPyV VP1, 41.6 kDa) forms spherical oligomers – VLPs – containing up to 360 monomers as described previously (Norkiene *et al*., 2015a). MC polyomavirus recombinant major capsid protein VP1 (MCPyV VP1, 46.6 kDa) forms spherical oligomers – VLPs, containing up to 360 monomers as described previously (Norkiene *et al*., 2015a). Measles virus recombinant nucleocapsid protein (MeV N, 58.0 kDa) forms filamentous structures as described previously (Samuel *et al*., 2003; Zvirbliene *et al*., 2007). Mumps virus recombinant nucleocapsid protein (MuV N, 66 kDa) also forms filamentous structures as described previously (Samuel *et al*., 2002a). All viral proteins were expressed in yeast expression system and purified by CsCl density gradient centrifugation (Norkiene *et al*., 2015a; Samuel *et al*., 2002a; Samuel *et al*., 2003; Slibinskas *et al*., 2004).

### VLP production, purification and analysis

Purification of VLPs of recombinant PyV VP1 proteins and electron microscopy were carried out as described previously (Norkiene *et al*., 2015b). Briefly, *S.cerevisiae* yeast biomass after the induction of recombinant proteins synthesis, were mechanically homogenized in DB450 buffer (450 mM NaCl, 1 mM CaCl_2_, 0.25 M L-Arginine and 0.001% Trition x-100 in 10 mM Tris/HCl-buffer, pH 7.2) with 2 mM PMSF and EDTA-free Complete Protease Inhibitors Cocktail tablets (Roche Diagnostics, Mannheim, Germany), and its supernatant was transferred onto 30-60% sucrose gradient. After overnight centrifugation (at 4 °C) at 100,000×g (Beckman Coulter Optima L-90 ultracentrifuge) collected 2 mL fractions were analysed by SDS-PAGE. The mixture of fractions containing PyV VP1 proteins diluted in DB150 buffer (150 mM NaCl, 1 mM CaCl_2_, 0.25 M L-Arginine and 0.001% Trition x-100 in 10 mM Tris/HCl-buffer, pH 7.2) and VLPs were concentrated by ultracentrifugation at 100,000×g for 4 h (at 4 °C). Thereafter, pellets containing VP1 were subjected to ultracentrifugation overnight on CsCl gradient (1.23-1.46 g/mL density) at 4 °C. One millilitre fractions of formed gradient were collected and analysed by SDS-PAGE. Positive fractions were pooled, diluted in DB150 buffer and concentrated as described above. The isolated VLPs were dissolved in PBS, dialysed and stored in PBS with 50% glycerol. The VLP formation was verified by examination of the purified proteins using Morgagni-268 electron microscope (FEI, Inc., Hillsboro, OR, USA). The protein samples were placed on 400-mesh carbon-coated palladium grids (Agar Scientific, Stansded, UK) and stained with 2% aqueous uranyl acetate.

### Cell cytotoxicity assays

Cell cytotoxicity was measured using lactate dehydrogenase (LDH) release assay (LDH cytotoxicity detection Kit). A quantity of 50 μL of cell supernatants was used to perform the cytotoxicity assay according to the manufacturer’s protocol. Briefly, 50 μL of supernatant was mixed with freshly prepared LDH reagent and incubated at 37 °C. After 30 min absorbance was measured at 490 nm using Multiskan GO microplate spectrophotometer (Thermo Fisher Scientific Oy, Finland).

To determine cell viability PI/Hoechst nuclear staining was also used. Nuclei were stained with 1.25 μg/ml PI and 1 μg/ml Hoechst33342 in cell culture medium for 30 min. The cells were washed with PBS. The fluorescent signal was measured by taking photos automatically with fluorescence microscope EVOS FL Auto. Images were taken using a 20× objective. Viability was quantified according to a ratio of PI (dead cells) and Hoechst (all cells), expressed in percentages.

### Quantitation of cytokines in cell culture supernatants

ELISA kits for the measurement of human cytokine – IL-6, IL-10, IL-1β and TNF-α – levels in cell culture supernatants were used (#88-7066, #88-7106, #88-7261, #88-7346, Thermo Fisher Scientific, USA). ELISA kits are based on the sandwich immunoassay technique. Supernatants were used diluted up to 1:600. All procedures were performed according to manufacturers’ protocols. In the last step 3,3’,5,5’-tetramethylbenzidine (TMB) substrate solution was added to each well. The plates were monitored for 15 min for colour development, the reaction in wells was stopped with 3.6% H_2_SO_4_ solution and the wells were read at 450 nm with reference wavelength at 620 nm using Multiskan GO microplate spectrophotometer. A standard curve was generated from cytokine standard and the cytokine concentration in the samples was calculated.

### Detection of ASC speck formation in THP-1 macrophages

THP-1 monocytes expressing ASC fused with green fluorescent protein (GFP) was used for ASC speck formation. The cells were cultivated in RPMI 1640 medium supplemented with 10% FBS, 1% P/S, 100 μg/mL normocin and 100 μg/mL selective antibiotic zeocin. THP-1-ASC-GFP cells were differentiated to macrophages as origin THP-1 using RPMI 1640 medium supplemented with 10% FBS, 1% P/S and PMA. THP-1-ASC-GFP macrophages were treated with viral proteins in serum-free RPMI. Thirty minutes before the treatment termination, Hoechst33342 was added to stain cell nuclei. After 24 h of treatment cell culture medium was replaced to FluoroBrite DMEM. ASC specks were analysed with EVOS FL Auto fluorescent microscope (Life Technologies, USA) by taking photos with 20× objective. Cells were counted according to the number of nuclei. ASC speck number per cell was counted using image processing program ImageJ.

### Measurement of active caspase-1

Active caspase-1 was detected using Fluorescent Labeled Inhibitors of Caspases (FLICA) assay according to manufacturer’s protocol. Briefly, FLICA reagent (FAM-YVAD-FMK – caspase-1 inhibitor probe) was added after the 15 h treatment with viral proteins and incubated for 1 h. Cells were washed three times and stained with Hoechst33342 at 1 μg/ml and PI at 1.25 μg/mL. After washing cells were analysed directly by fluorescence microscope EVOS FL Auto using a 20× or 40× objective.

### SDS-PAGE and Western blot analysis

To determine cleaved caspase-1 p20 fragment Western-blot assay (WB) was performed. After cell treatment with viral proteins, supernatant was collected and centrifuged at 600 × g for 10 min to remove cellular debris. Then, the protein content of the supernatants was concentrated 10x using centrifugal filters with 10 kDa cutoff (#UFC501096, Amicon, Merck). The concentrated samples were boiled in a reducing sample buffer and separated in 4-12% polyacrylamide gel (#NW04122BOX, Thermofisher Scientific) electrophoresis (PAGE) in MES SDS running buffer (#B0002 Invitrogen, Thermofisher Scientific). The proteins from the SDS-PAGE gel were blotted onto 0.2 µm nitrocellulose (NC) membrane (#LC2000, Invitrogen, Thermofisher Scientific) by wet transfer. The membrane was blocked with 5% BSA in PBS for 1 h at RT and rinsed with TBST. The membrane was then incubated with primary antibodies in TBST with 1% BSA overnight at 4 °C. The primary antibodies against human caspase-1 (clone Bally-1, #AG-20B-0048-C100, Adipogene) were used at 1:1000 dilution. Thereafter, the membrane was incubated with secondary antibodies Goat Anti-Mouse IgG (H+L)-HRP Conjugate (Bio-Rad) diluted 1:5000 in TBST with 1% BSA for 1 h at RT. The horseradish peroxidase (HRP) enzymatic reaction was developed using chemiluminescent substrate (#34094, Thermofisher Scientific).

### Immunocytochemistry for studying the uptake of VLPs by macrophages

Cells were stained in IbidiTreat 96 well µ-plates. After the treatment cells were washed with PBS and fixed in 4% PFA dissolved in PBS for 15 min and permeabilised with 0.1% Triton X**-**100 prepared in PBS for 10 min. Blocking solution – PBS containing 2% BSA was applied for 30 min followed by two washing steps. The primary antibodies rabbit polyclonal anti-CD68 (1:100; #25747-1-AP, Proteintech) and mouse anti-PyV VP1 VLPs (monoclonal antibodies of hybridoma supernatant at dilution 1:2) were added to the blocking solution and incubated overnight. The following secondary antibodies were used respectively: goat anti**-**rabbit (1:1000) and goat anti**-**mouse (1:1000). The secondary antibodies were applied for 2 h followed by two washing steps. Hoechst33342 was used for nuclear staining at 1 μg/mL for 30 min in PBS. The images were taken using a 40× objective. CD68 was used as a macrophage and lysosomal marker. The experiment was imaged using EVOS FL Auto fluorescence microscope (ThermoFisher Scientific, USA). Acquired images were processed using ImageJ (Wayne Rusband; National Institute of Health, Bethesda, MD, USA).

For the immunocytochemistry, in-house generated murine MAbs against recombinant NLPs and VLPs were used (#MAb clone – virus antigen indicated): #7C11 – MeV N (Zvirbliene *et al*., 2007); #5E3 – MuV N (Samuel *et al*, 2002b); #5G8 – KIPyV VP1; #11A2 – MCPyV VP1.

### Microscopy

All images were taken with fluorescence microscope EVOS FL Auto (#AMAFD1000, Thermofisher Scientific). EVOS imaging systems use LED light cubes, which combine bright LED illumination with excitation and emission filters into single components. PI signal was detected in RFP light cube; Hoechst33342 signal – in DAPI light cube (AMEP#4650); GFP and AlexaFluor 488 signal – in GFP light cube (AMEP#4651); PI signal – in RFP light cube (AMEP#4652); AlexaFluor 594 signal – in TxRed light cube (AMEP#4655). The following objective lenses were used 20× (#AMEP4682) and 40× (#AMEP4683). Acquisition software EVOS® FL Auto v1.6 was used. Images were prepared with Image J program. Firstly, appropriate pseudo-colour was added at images of 8-bit format. Then, the images were converted to RGB colour format. Finally, brightness and contrast were adjusted equally to all images per channel. Images (fluorescent signal and object counting) were analysed with ImageJ. The final figures were arranged using Adobe Photoshop without any brightness/contrast and colour manipulations.

### Cell preparation for single cell RNA sequencing

THP-1 macrophages for RNA sequencing analysis were treated for 15 h with PyV- derived VLPs (20 µg/ml). After the treatment cells were washed with PBS and detached with TrypLE reagent after 15 min incubation at cell culture incubator (37 °C, 5% CO_2_). After centrifugation cells were washed twice with 1x RNase-free PBS (cat#AM9624, Invitrogen). After final centrifugation cells were resuspended in RNase-free PBS and kept on ice till cell preparation for RNA sequencing analysis

### Single-cell RNA sequencing

Single-cell RNA sequencing (scRNAseq) was performed using a modified version of the inDrops method (Klein *et al*, 2015; Zilionis *et al*, 2017) While the original version is based on the CEL-Seq protocol (Hashimshony *et al*, 2012) that involves linear cDNA amplification by *in vitro* transcription, the modified version relies on template switching and cDNA amplification by PCR, as in the Smart-seq protocol (Ramsköld *et al*, 2012). First, single cell transcriptomes were barcoded in 1-nl droplets by co-encapsulating: i) barcoding hydrogel beads; ii) the reverse-transcription/lysis (RT/lysis) mix; and iii) the cell suspension. Cell encapsulation was performed using the microfluidic device described in (Zilionis *et al*., 2017) on an Onyx platform (Droplet Genomics). Maxima H- minus (ThermoFisher, cat. no. EP0751) was used for reverse-transcription with template switching. RT was performed for 1h at 42 °C followed by heat inactivation for 5 min at 85 °C. The emulsion was then broken and the pooled material was taken through library preparation for Illumina sequencing, which involves the following steps: i) cDNA amplification (Terra PCR direct Polymerase, Takara, cat. no. 639270), ii) fragmentation and adapter ligation (NEBNext® UltraTM II FS DNA Library Prep Kit for Illumina, NEB, cat. no. E7805S), and iii) indexing PCR (KAPA HiFi HotStart ReadyMix PCR Kit, Roche, cat. no. KK2601). Sequencing was performed on a single Illumina NextSeq run (NextSeq 500/550 High Output Kit v2.5 (75 Cycles), Illumina, cat. no. 20024906).

### ScRNAseq raw data processing

The <u>solo-in-drops</u> pipeline (https://github.com/jsimonas/solo-in-drops), which is a wrapper around <u>STARsolo</u> (https://github.com/alexdobin/STAR/blob/master/docs/STARsolo.md), was used for obtain cells x genes expression matrices. STARsolo expects as input two fastq files per library, one containing the barcode information and the other the transcript information. Meanwhile, the custom scRNAseq protocol used in the current study outputs 3 fastq files per library: barcode half 1, barcode half 2, and transcript. Solo-in-drops prepares the data for compatibility with STARsolo. STAR (version 2.7.6a) was run with the following parameters: -- soloType CB_UMI_Simple, -- soloUMIfiltering MultiGeneUMI, -- soloCBmatchWLtype 1MM_multi_pseudocounts. Homo sapiens (human) genome assembly GRCh38 (hg38) was used as the reference.

### ScRNAseq count data analysis

Jupyter notebooks with commented code for scRNAseq data analyses, including data filtering, normalization, visualization, clustering, cell population annotation, differential gene expression (DGE) analysis, cell cycle scoring, and plots used in Figures 9-11 and EV9-11 are provided on GitHub (github.com/rapolaszilionis/Luciunaite_et_al_2022). The Scanpy toolbox was used for data analysis. Transcriptomes with fewer than 900 total counts were excluded. No filtering on the fraction of mitochondrial counts was performed. UMAP (Becht *et al*, 2018) was used for data visualization in 2D. Leiden clustering was used to divide the graph into clusters, which were annotated based on their gene expression profiles. Gene Ontology (GO) gene set enrichment analysis (Fig. 9D, 10E) was performed using the tool available online (http://geneontology.org) using Panther GO-Slim Biological Processes as the annotation data set. For DGE analyses (Figs. 9C, 10E), the Mann-Whitney U test was used to test for significance, and the Benjamini-Hochberg procedure was used to correct for multiple hypothesis testing.

### Statistical analysis

All statistical analyses were performed with GraphPad Prism 9.2.0 (GraphPad Software, Inc., La Jolla, CA). The data in the figures are represented as individual data points from at least 6 independent experiments using box plots (showing minimum, first quartile, median, third quartile and maximum) or bar graphs. Independent experiments referred to as N means the number of independent cell culture preparations and n means the number of technical repeats. Normality test was carried out to test if the values come from a Gaussian distribution. Statistical comparisons of vehicle controls versus treatment were performed with one-way ANOVA in conjunction with a Tukey’s multiple comparison test or Student’s t-test. A Kruskal–Wallis test with Dunn’s post hoc test was used for non-parametric data. Differences with p value less than 0.05 were considered to be statistically significant: *p < 0.05, **p<0.01, ***p < 0.001, ****p < 0.0001. To emphasise non-significant results ns was used.

## Acknowledgements

This study was supported by the Research Council of Lithuania, grant No. S-SEN-20-11. We would like to thank prof. L. Mažutis for cell line THP-1.

## Author Contributions

Asta Lučiūnaitė designed experiments, analysed the data and wrote the manuscript. Indrė Dalgėdienė and Kristina Mašalaitė contributed to the experiments and manuscript writing. Milda Norkienė was responsible for VLP preparation. Andrius Šinkūnas performed single-cell RNA sequencing. Rapolas Žilionis was responsible for single-cell RNA sequencing analysis. Indrė Kučinskaitė-Kodzė and Alma Gedvilaitė helped to design the experiments and were responsible for manuscript revision. Aurelija Žvirblienė was responsible for the final manuscript revision. All of the authors reviewed the manuscript and approved its final version.

## Conflicts of Interest

R.Ž. and A.Š. are employed at Droplet Genomics. R.Z. is a shareholder at Droplet Genomics. These commercial relationships are unrelated to the current study. All authors declare no conflict of interest regarding the publication of this paper.

## Data Availability

Single-cell RNAseq-related datasets and computer code produced in this study are available in the following databases:

- Fastq files and count matricesGene Expression Omnibus GSEXXXXX (https://www.ncbi.nlm.nih.gov/geo/query/acc.cgi?acc=GSEXXXXX)
- Code for analyses and figures: GitHub (https://github.com/rapolaszilionis/Luciunaite_et_al_2022)

All other data used to support the findings of this study are included within the article.

## Tables and their legends

Only expanded view tables uploaded as a separate file.

**Table EV1. Bulk-like scRNAseq data differential gene expression analysis results**

**Table EV2. GO gene set enrichment analysis results for genes commonly enriched in VLP treated samples**

**Table EV3. Data underlying** **Fig 10E**

## Expanded View Figure legends

**Figure EV9.**
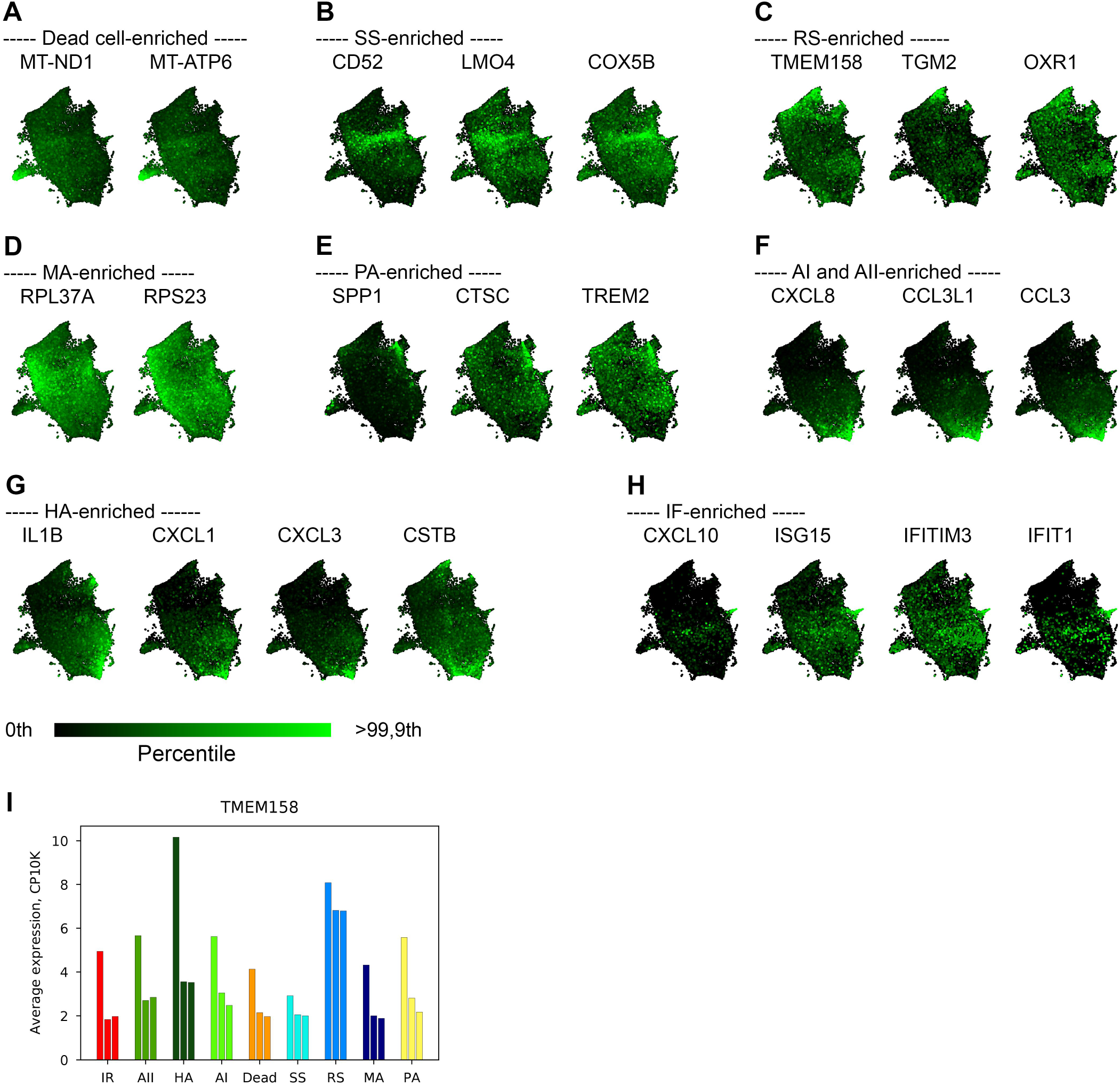
Bulk-like scRNAseq data analysis. Volcano plot showing the result of a bulk-like comparison of MCPyV vs KIPyV VLPs. Differentially expressed genes (DGEs) were defined as having an absolute fold-change >1.5 and FDR < 0.05 (Mann-Whitney U test).

**Figure EV10.**
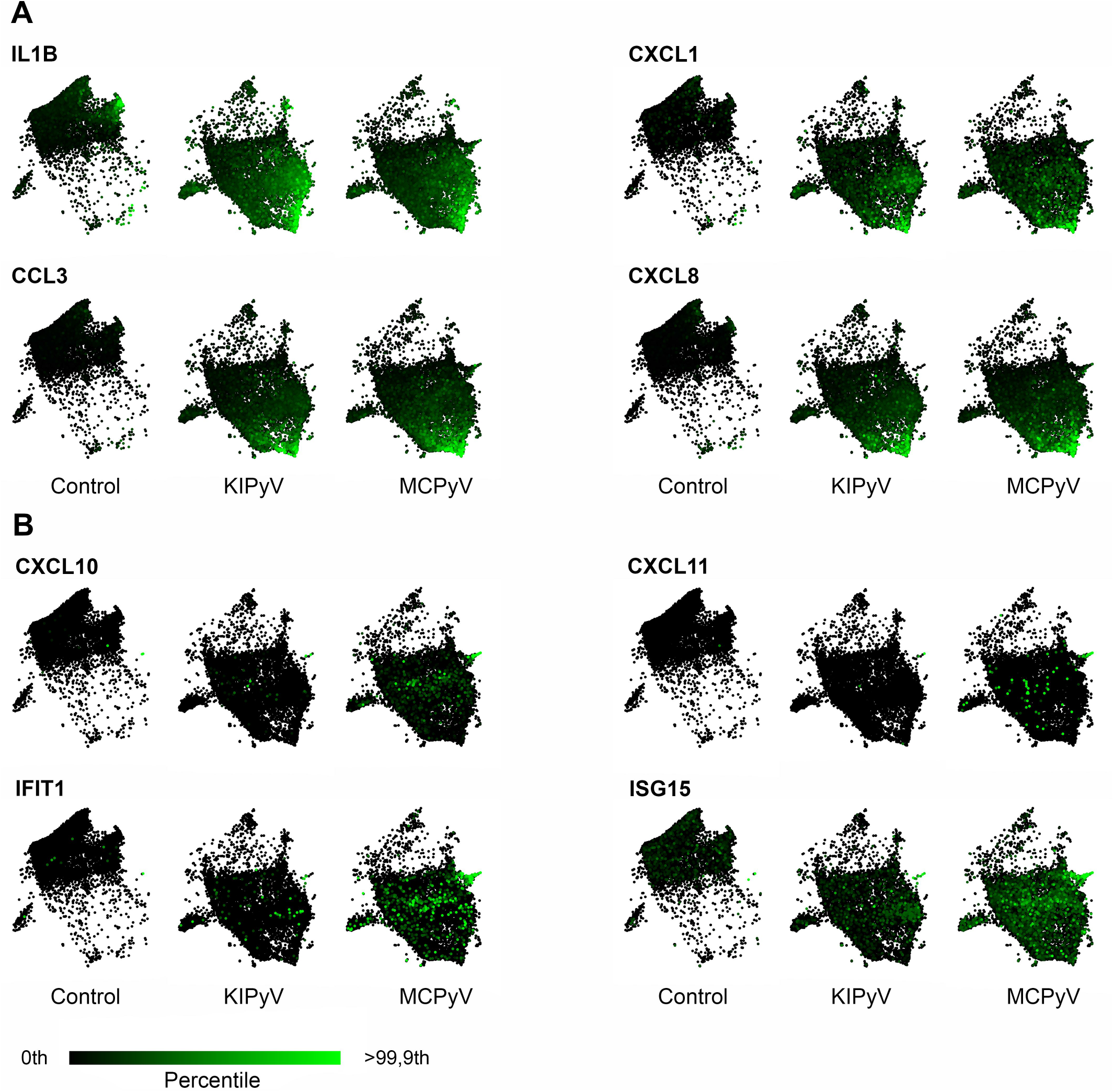
UMAP plots combining all conditions coloured by expression of selected genes. A-H Plots of enriched genes in separate populations are shown: (A) dead cells, (B) SS population, (C) RS population, (D) MA population, (E) PA population, (F) HA population, (G) AI and AII populations, and (H) IF population. I Bar chart of average *TMEM158* expression in individual populations and conditions. Condition order left-to-right: Control, KIPyV VLPs, MCPyV VLPs.

**Figure EV11.**
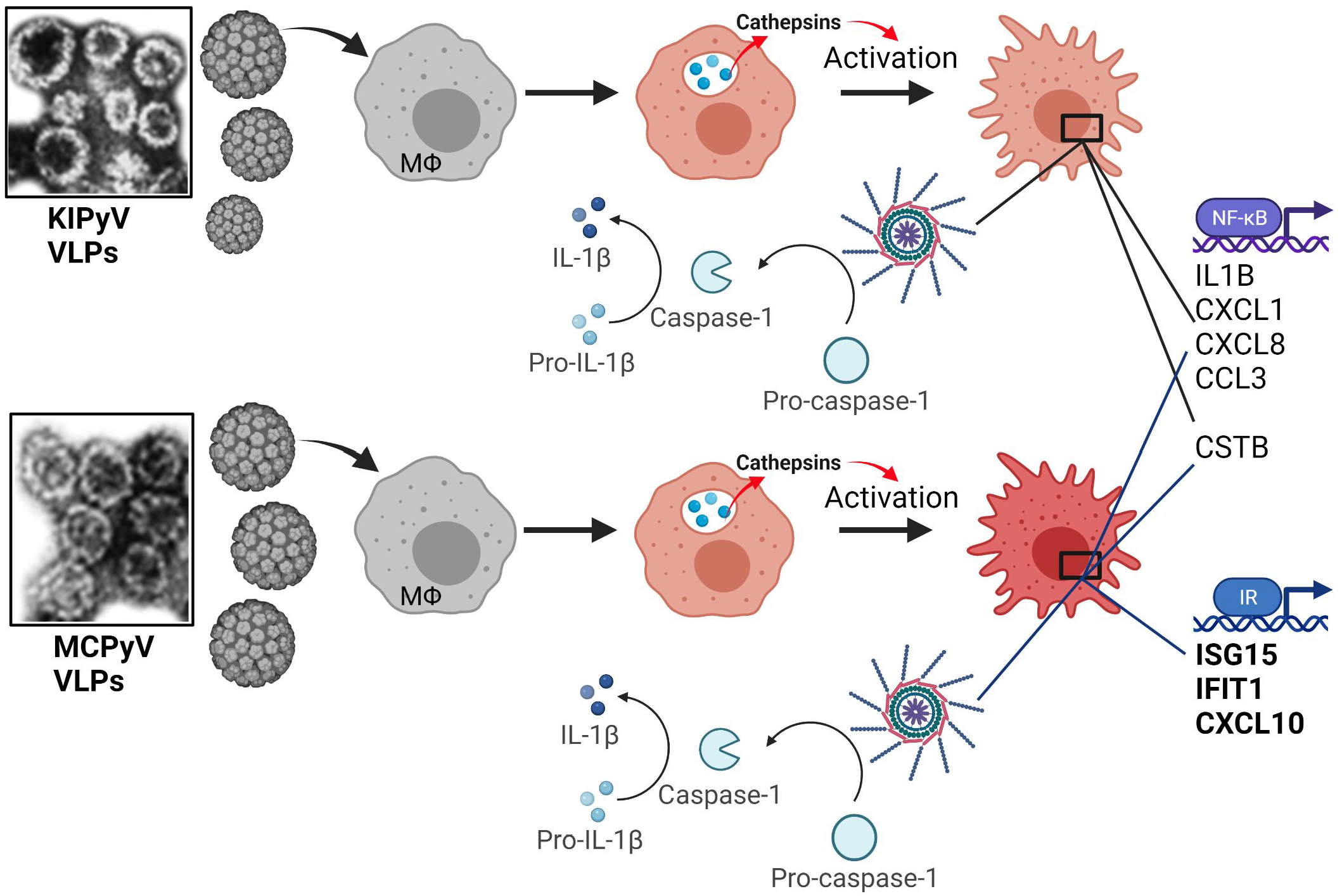
Examples of genes enriched in different treatments – control, KIPyV and MCPyV. A, B UMAP plots colored by expression of selected genes in each condition separately. For a given gene, the 3 plots are saturated at the same absolute value to allow comparison. *IL1B*, *CXCL1*, *CCL3*, and *CXCL8* are examples of genes upregulated in both KIPyV and MCPyV conditions relatively to the control (A). *CXCL10*, *CXCL11*, *IFIT1*, and *ISG15* are expressed in the MCPyV-specific IR population (B).

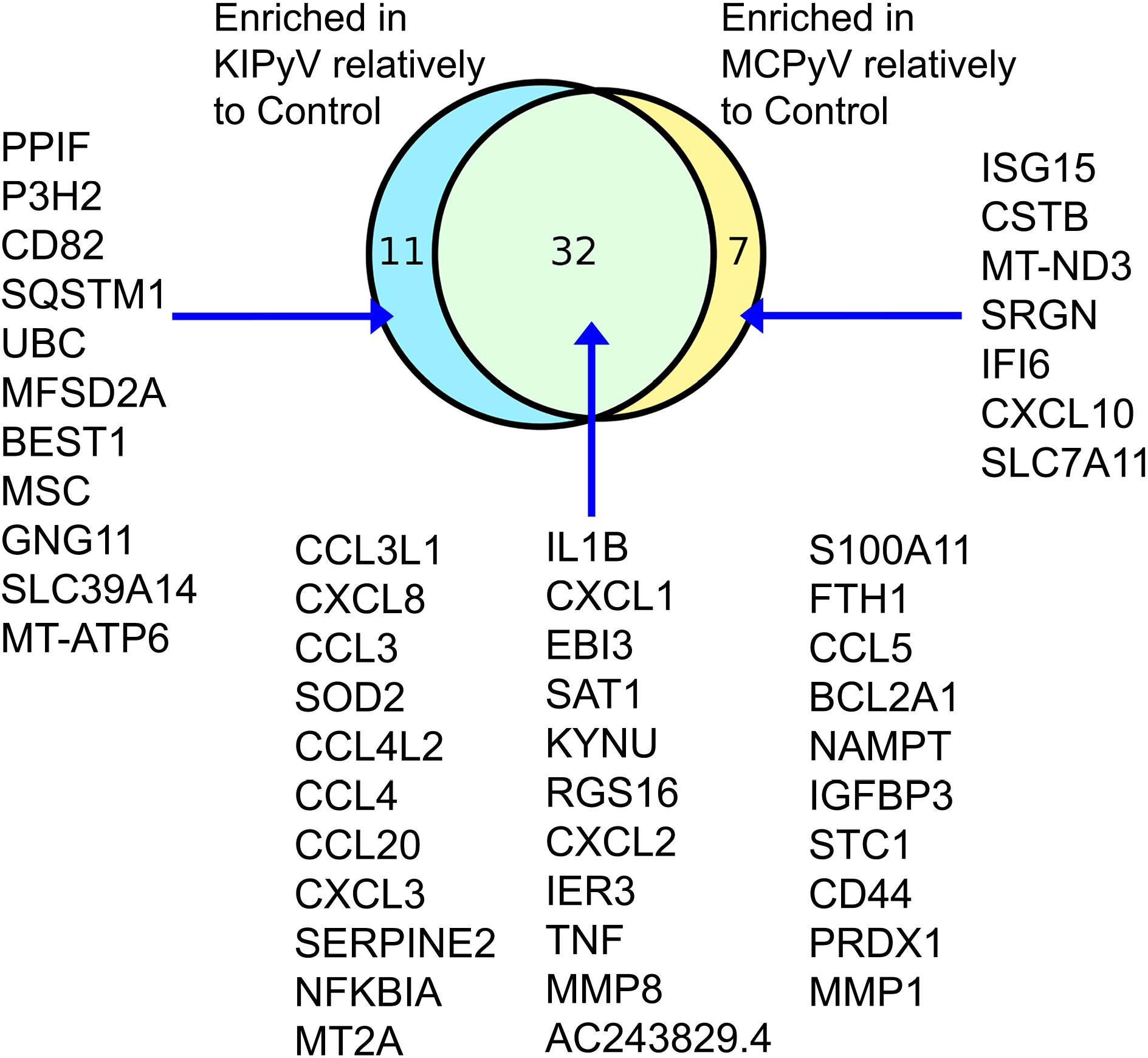

